# A mammalian fibroblast zoo reveals barriers to cross-species virus replication and differential regulation of antiviral immunity

**DOI:** 10.64898/2026.11.04.763981

**Authors:** Clayton K. Mickelson, Mark Salnikov, Frances K. Shepherd, Lauren M. Pross, Georgia McClain, Swechha Mainali Pokharel, H. Heinrich Hoffmann, Timo Schoon, Girish Pulinkala, Nakarin Pamornchainavakul, Kayla Hoffman, Cera A. McDonald, Esther Vogt, Tristan T. Neal, Adam J. Gilbertsen, Dira S. Putri, Jordan T. Becker, Mark J. Pierson, Yue Li, Matthew T. Aliota, Kenneth H. Dinnon, Stephanie Zec, Susie Garity, Anne Rivas, Jim Rasmussen, Arno Wuenschmann, Michael Gale, Charles M. Rice, Jan Komorowski, Kimberly VanderWaal, Sara Cherry, Ryan A. Langlois

## Abstract

Virus zoonosis represents a continual threat to humans. Viruses emerge in humans directly or indirectly through interactions with mammals. However, our understanding of virus-host susceptibility across the diversity of mammals is limited and experimental model systems are restricted to a very small set of model organisms. Mammals are highly diverse and different habitats, lifestyles, diets, and exposures to different pathogens have driven substantial genetic diversity. We generated a ‘library’ of primary fibroblasts across 86 mammalian species and used these cells to build an atlas of virus-host susceptibility across a panel of arboviruses that infect diverse mammals including humans. Perturbing the innate immune system revealed some virus-host pairs where restriction is controlled by the immune system and others where there is an incompatibility with host factors. Transcriptomic profiling of the library in response to virus sensing mimicked by poly(I:C) or antiviral signaling with type I interferon reveals a core set of conserved genes that respond as well as substantial species- and order-specific responses. Together these data provide a comparative atlas of innate immune responses across mammals and offer a window into how these systems have evolved across animals.

## Introduction

Viruses that enter the human population from zoonotic reservoirs can have devastating consequences. Over the last century, there have been several significant zoonotic events including 1918 influenza A virus, human immunodeficiency virus, Ebola virus, Zika virus, and SARS-CoV-2, resulting in over 100 million deaths(*1, 2*). There is also continual spread of viruses into new areas. Indeed, viruses that previously did not cause significant outbreaks are infecting humans at a rate that is the highest in human history, and this is predicted to increase for several reasons. For example, climate change and urbanization are forcing species into new habitats, resulting in new cross-species interactions, further increasing the odds of pathogen spillover into new hosts(*3*). Arthropod-borne infections are particularly concerning sources of potential spillover. Rising temperatures and altered rainfall patterns are expanding the habitats of virus vectors into new regions while urbanization is expanding the breeding grounds for mosquitoes(*4, 5*). This increases the odds of exposure to new species. Indeed, 2024 stands as the worst year on record for dengue virus, shattering previous global records with over 14 million reported infections(*6*). Arboviruses are emerging and re-emerging globally and there is a lack of off-the-shelf antiviral therapeutics, vaccines, and large-scale diagnostic testing capacity for most arboviruses. While arboviruses must infect a blood-feeding arthropod and a vertebrate to be maintained in nature, we have a poor understanding of which vertebrate hosts are susceptible within these expanding geographic locations. Arboviruses are highly diverse with both positive and negative sense genomes and divergent replication and innate immune evasion strategies. The study of virus-host interaction has focused on humans and a few model organisms, leaving significant gaps in our understanding of molecular interactions that may drive resistance and susceptibility in diverse species, some of which may serve as reservoirs or may become infected in the future due to spillover. Understanding barriers to infection will reveal new mechanisms for potential interventions.

Viruses face several barriers when encountering new species. First, as obligate intracellular pathogens, viruses must hijack host factors to complete their life cycle. Therefore, they require compatible entry receptors and host factors for downstream activities and successful infection. Variation in these factors can have striking effects on susceptibility; for example, an insertion in the entry receptor MXRA8 makes cows resistant to chikungunya virus (CHIKV) infection(*7*). The spectrum of permissive and non-permissive hosts is not understood, impacting our ability to predict spillovers. Second, successful viruses must evade innate immune antiviral responses to establish infection. Viruses are recognized by host pattern recognition receptors leading to the production of type I interferon (IFN). IFN then signals in an autocrine and paracrine manner to induce hundreds of interferon-stimulated genes (ISGs) which have antiviral activity thereby blocking viral infection. Viruses evade these responses by encoding antagonists that can block virus sensing and/or these antiviral effectors. Thus, there is a molecular arms race between viruses and hosts which has shaped both of their evolutionary trajectories. Both pro- and antiviral host factors are under heavy selection and are highly divergent across species(*8–11*). Comparative studies have uncovered novel mechanisms of virus resistance (*12, 13*) but have been restricted to only a few model and non-model species and often use only single genes expressed in human cells, limiting our understanding.

Here we built and characterized a panel of primary dermal fibroblasts from diverse mammal species. We used this fibroblast zoo to interrogate resistance and susceptibility to arboviruses in the three groups of arboviruses that infect humans: Togaviridae, Flaviviridae and Bunyaviricetes. Fibroblasts represent a relevant target for arboviruses at the site of infection when arthropods take a blood meal through the skin and infect these cells(*14*). Fibroblasts are readily obtainable from dermal biopsies and straightforward to expand permitting large-scale experimentation. We validated cells from 86 species and challenged them with 18 viruses to determine baseline infection susceptibility using automated microscopy and image analysis. We evaluated pro- and antiviral immune barriers by blocking JAK-STAT signaling. Species where infection increases in the presence of a JAK inhibitor represent those virus-host pairs where IFN restricts infection.

Failure to infect even with immune blockade suggests incompatibility with proviral host factors. For species that are highly permissive, we assessed whether this was due to antagonism of IFN signaling by pretreating with IFN and monitoring infection. This revealed several previously unidentified situations where viruses could evade antiviral immunity and replicate to high levels. Altogether these studies generated 15,867 pairwise comparisons creating the largest experimental eukaryotic virus-host infection compatibility dataset ever assembled. We complemented these studies with transcriptomics defining the innate immune landscape across 63 diverse species. This revealed both conserved responses and species-, family-, and order-specific responses highlighting the diversity of innate immunity across mammals. Finally, we used machine learning to uncover virus and host features that impact susceptibility and resistance. Together this manuscript provides an atlas of virus-host interactions and dynamic transcriptomes, which can be further mined to uncover novel mechanisms of virus resistance and susceptibility across mammals.

### Differential susceptibility across viruses and hosts

To determine how evolutionary divergence impacts susceptibility to infection, we built and characterized a panel of primary dermal fibroblasts from tissue donated from traditional and non-traditional animal model researchers, zoos, veterinary diagnostic laboratories, and departments of natural resources across the US using a standardized protocol within a single laboratory (Table S1). Fibroblasts are readily obtainable from dermal biopsies and straightforward to expand, permitting large-scale experimentation. We generated primary fibroblasts from 86 species including 14 of the 26 orders in Mammalia with multiple genetically diverse individuals from 42 species (Fig. 1A and Supplementary Table 1). We used this fibroblast zoo to interrogate resistance and susceptibility to arboviruses in the three groups of arboviruses that infect humans: Togaviridae, Flaviviridae, and Bunyaviricetes. Fibroblasts represent a relevant target for arboviruses at the site of infection when arthropods take a blood meal through the skin(*14*). Furthermore, primary cells have significant advantages over transformed cells which often have mutations in cell cycle and innate immune pathways, impacting interpretation of virus-host interaction studies(*15*). We screened cells from 81 species with a panel of 18 arboviruses which generated 1443 pairwise observations and revealed a unique fingerprint of infectivity of viruses across hosts (Fig. 1B). Importantly, many infection profiles mirror what is seen in nature. For example, CHIKV is known to infect humans and many rodent species but is restricted in cows(*7*), which was reflected in our *in vitro* infection data. Additionally, Zika virus exhibited low infectivity across all primates but was completely restricted in the new world bat species tested(*16*). Additionally, this model allows for evaluation of virus host pairs that currently do not interact but could in the future (e.g. Rift Vally fever virus (RVFV) in north American carnivores). Principal components analysis revealed that many related host species shared similar infection profiles (Fig. 1C) suggesting phylogenetic relatedness is a major contributor to infection potential. However, there are many examples where species did not cluster with other order-level representatives, such as alpaca, golden hamster, and common marmoset, suggesting that genetic divergence within related species can control susceptibility.

**Figure 1.**
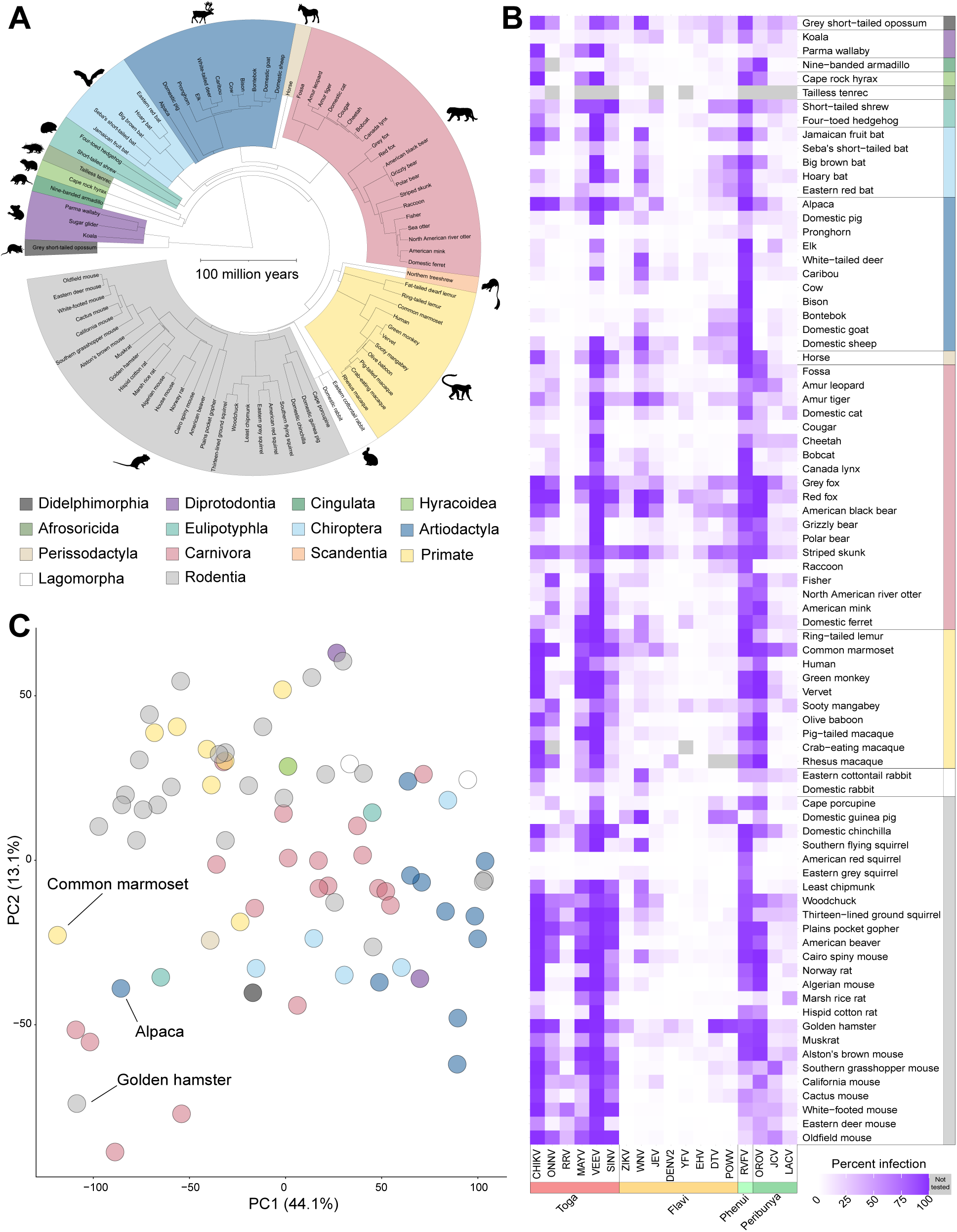
Differential susceptibility across viruses and hosts. (A) Phylogenetic tree of 86 mammalian species used in this study. Branch distance represents years of divergence. (B) 81 species were infected with the indicated 18 viruses and % infection determined to define susceptibility. See Table S2 for MOI, time point, and strain for each virus. (C) Principal components analysis of % infectivity data from B with individual species colored by order.

The virus-by-host matrix provides a snapshot of infection susceptibility. Infectivity can be dictated by a number of experimental factors including infectious dose, time point, and detection methods. Therefore, we evaluated whether these technical differences were driving phenotypes. First, we infected with several multiplicities of infection (MOI) and time points resulting in 5724 total pairwise comparisons (Supplementary Fig. 1A). As expected, we found that infectivity increased with increasing MOI and time (Supplementary Fig. 1B, C). Importantly, there was still a full dynamic range of susceptibility at late time points and high MOI. Second, we evaluated viral antigen using a second readout, virus-derived dsRNA and found concordant infection phenotypes (Supplementary Fig. 1D). This suggests that the experimental data captures a representative range of infectivity outcomes across diverse species. Moreover, it is possible that heterogeneity across individuals, batches or plating density drives differences we observe in susceptibility across species. We observe strong concordance in infectivity across multiple independent experiments, genetically outbred individuals, and closely related virus strains (Supplementary Fig. 1E, F, and G). Together these data demonstrate that the fibroblast zoo can be used to characterize infection profiles across diverse viruses and hosts with strong reproducibility of major infection phenotypes.

### Innate immunity presents a major barrier to infection across species

Viruses confront several barriers when infecting new species. They must utilize host factors to enter cells and replicate their genomes while also evading antiviral immune responses. We distinguished whether poor infection in species was due to a defect in replication or a defect in IFN signaling by blocking JAK-STAT signaling. Species where infection increases in the presence of a JAK inhibitor represent those virus-host pairs where IFN produced restricts infection. We utilized the JAK inhibitor ruxolitinib (JAKi) because it targets a highly conserved pocket in the active site in JAK1/2 and functions across mammals, birds, and fish to block IFN signaling(*17, 18*). This treatment revealed two major classes of virus-host groups: those that remain refractory to infection and those where we observe an increase in infection compared to baseline (Fig. 2A). We reasoned that there may be cases where increased cytopathic effect would skew the analysis of how JAK inhibition impacted infectivity, therefore we evaluated cell counts and found that they did not change dramatically despite the absence of JAK/STAT signaling and higher infection (Supplementary Fig 2A). Interestingly, there were a few cases where cytopathic effect was markedly higher (e.g. skunks during Powassan infection) which may indicate different degrees of susceptibility and pathology (Supplementary Fig 2A). Virus-host pairs with failed replication even in the presence of JAKi likely represent incompatibility with proviral factors (e.g. entry receptors, RNA-dependent RNA polymerase cofactors, Golgi trafficking proteins, etc.). Phylogenetic patterns of resistance were apparent, including dengue virus which failed to infect all bat species tested and yellow fever virus which failed in rodents. These examples suggest an evolutionarily conserved lack of host factor expression and/or sequence divergence of the host factor among certain phylogenetic lineages.

**Figure 2.**
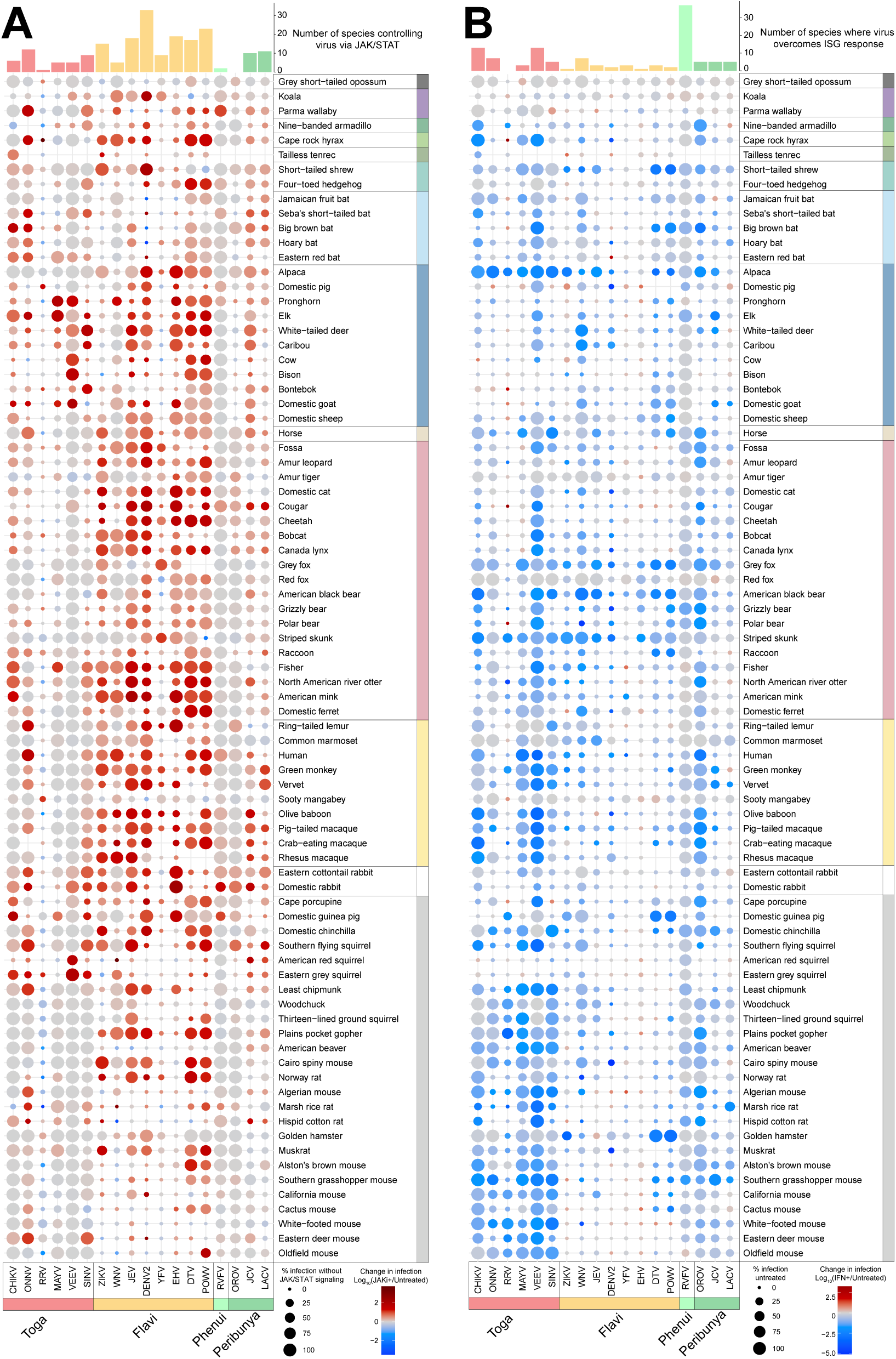
Innate immunity is a barrier to replication across many virus-species pairs. Cells from 81 species were pretreated for 1hr with ruxolitinib (JAKi) (A) or for 2h with 1000 U/mL of uIFN (B) or with media alone (untreated) and were analyzed 24-72h later by immunostaining with virus-specific antigen. (A) Percent infection with JAKi (size of dot) was determined by imaging. Log10 fold change in infection of JAKi-treated cells over untreated cells is shown as color of the dot (rescue of infection in red). Bar graph at the top represents the number of mammalian species which can control the virus via JAK/STAT signaling. A mammalian species was considered to control a virus via JAK/STAT signaling if the percent infection in untreated cells was less than or equal to 5% and the infection in JAKi-treated cells was greater than 5% and there was at least a 0.7 log10 fold change between the JAKi-treated and untreated percent infection values. (B) Percent infection of untreated cells (size of dot) was determined by imaging. Log10 fold change in infection of uIFN-treated cells over untreated cells is shown as color of the dot (repression of infection in blue). Bar graph at the top represents the number of mammalian species in which the virus can overcome the ISG response. A virus is considered to overcome the ISG response if the percent infection in untreated cells was greater than 5%, the percent infection in uIFN-treated cells was greater than 20%, and the log10 fold change between uIFN and untreated was between -0.21 and 0.5. See Table S2 for MOI, time point, and strain for each virus.

We next evaluated the number of species where replication is restricted by endogenous IFN signaling for each virus. We found that orthoflaviviruses were consistently under the strongest control by JAK/STAT-driven innate immunity (Fig. 2A). This is consistent with data demonstrating that some orthoflaviviruses have species-specific targeting of STAT1/2 and STING(*19–21*). Interestingly, several viruses have high infectivity across species at the MOIs used without JAKi. To determine if evasion of innate immune restriction was due to the infection dose used, we evaluated additional lower MOIs and additional time points (Supplementary Fig. 2B). In most cases, changing the MOI or time point only modestly impacted classification of innate immune restriction (Supplementary Fig. 2C). 22% of virus-host pairs presented with low susceptibility at baseline and upon JAK inhibition. These likely represent examples of host factor incompatibility barrier (Supplementary Fig. 2D).

Many viruses antagonize IFN signaling and thus JAK inhibition may not impact infection. However, treatment with IFN may inhibit infection due to the induction of antiviral ISGs. Therefore, we set out to determine the virus-host pairs that can be controlled by IFN. For these studies we pretreated cells with a mammalian “universal” IFNα (uIFN) and evaluated infectivity. We found that uIFN treatment reduced virus infection in at least one virus for all species except in marsupials, hedgehog, and ring-tailed lemur (Fig. 2B). These data suggest that uIFN signals in most species to sufficient levels to provide protection against infection. We found several examples of viruses able to replicate even in the face of IFN and antiviral gene expression, suggesting that they can evade antiviral effectors within select species (Fig. 2B). For example, Venezuelan equine encephalitis virus (VEEV), which has rodents (e.g. spiny mice and cotton rats) as a natural reservoir, readily infects many rodents and can be controlled by exogenous IFN. However, VEEV maintains high levels of replication in woodchuck and thirteen-lined ground squirrel in the presence of exogenous uIFN, which could be due to specialized innate immune evasion mechanisms. Conversely, RVFV, which encodes NSs that blocks transcription by degrading the highly conserved p62 subunit of the basal transcription factor TFIIH(*22, 23*), can overcome the antiviral response and replicate in many species pretreated with uIFN, demonstrating a generalist approach to innate immune antagonism across Mammalia.

### Entry is a species barrier to infection

It is well known that polymorphisms in entry receptors can pose significant barriers to infection. For example, previous sequence analysis of the CHIKV receptor Matrix Remodeling Associated 8 (MXRA8) identified an insertion in Bovinae species that was predicted and subsequently found to be defective for entry(*7*). As expected, CHIKV infected human and many other species of fibroblasts, while fibroblasts derived from cow and bison were resistant to CHIKV infection at 24 hpi (Fig 1B, 2A, 3A). We also observed low infection of CHIKV in eastern red bat and hoary bat during the first round of infection (Fig. 1B, 3A). However, when we allowed infection to proceed for 48h, we observed low levels of infection that were increased by JAKi treatment (Fig. 3B). Since JAKi treatment did not increase infection at 24h, these data suggested that there may be a block to infection early that is distinct from the later IFN-dependent block. To determine if the low infection at 24 hours was due to a defect in entry, we transduced eastern red bat and hoary bat fibroblasts with either vector control or murine MXRA8 (mMXRA8), which has been shown to be a functional receptor for CHIKV(*24*). Expression of mMXRA8 led to significant increases in infection of the eastern red bat at 24hpi with more modest increases in the hoary bat that was potentiated by JAKi treatment (Fig. 3C, Supplemental Fig. 3A). Live-cell imaging confirmed that expression of mMXRA8 led to rapid and complete infection of the eastern red bat, earlier than infection of human fibroblasts (Fig. 3D). In contrast, the hoary bat remains less permissive even at 48 hours upon IFN signaling blockade (Fig. 3D). Altogether, these studies reveal that some bats have evolved additional mechanisms to evade CHIKV infection.

**Figure 3.**
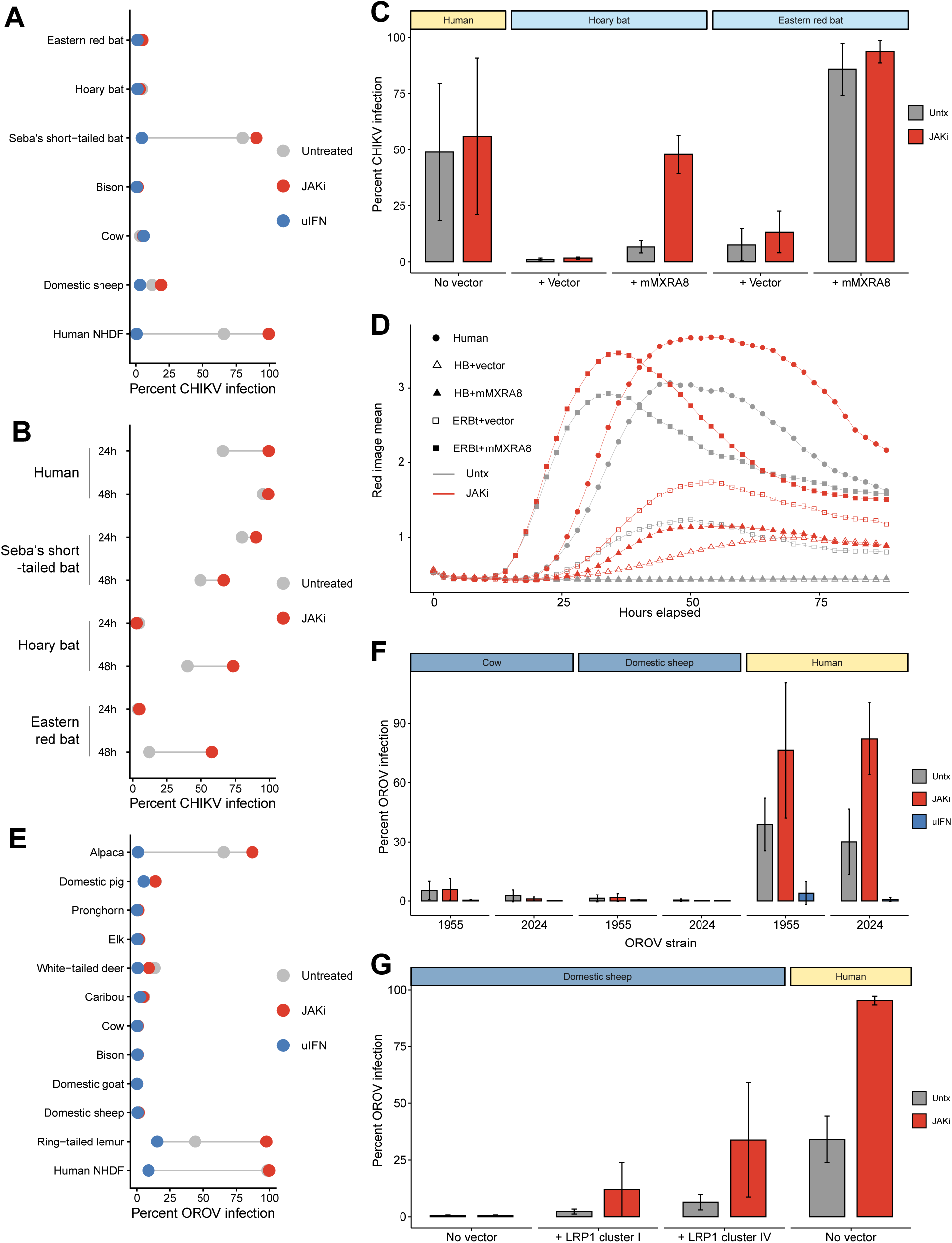
Viral entry is an important species barrier. (A) Indicated cells were treated with DMSO, JAKi, or uIFN and infected with CHIKV (MOI 0.2) and fixed and imaged at 24hpi (n = 1). (B) Indicated cells were treated with DMSO or JAKi, infected with CHIKV (MOI 0.2) and fixed and imaged at 24hpi or 48hpi (n = 1). (C) Indicated cells were transduced with murine MXRA8 (mMXRA8) or vector control and treated with DMSO or JAKi, infected with CHIKV (MOI 0.01 or 0.02, depending on experimental replicate) and fixed and imaged at 24hpi (n=2-5). (D) Indicated cells were transduced with mMXRA8 or vector control and were infected with CHIKV expressing mKate (MOI 0.01) and live-cell imaging was used to quantify infection with and without JAKi. HB = Hoary bat, ERBt = Eastern red bat. (E) Indicated cells were treated with DMSO, JAKi or uIFN and infected with OROV (1955 strain, MOI = 5) and fixed and imaged at 48hpi (human n=5, sheep n=2, all others n=1). (F) Cow, sheep and human cells were pretreated with DMSO, JAKi, or uIFN, infected with the 1955 or 2024 strain of OROV (MOI 5), and fixed and imaged at 48hpi (n = 3). (G) Sheep fibroblasts were transduced with human LRP1 cluster I or LRP1 cluster IV, selected cells were pretreated with DMSO or JAKi, or infected with OROV (1955) at an MOI of 5 and fixed 48 hpi (DMSO n=4, JAKi n=2).

We also observed that Oropouche virus (OROV) displayed very low to undetectable infection of many ruminants (Fig. 1B, 2A, and 3E). This is surprising as RVFV readily infects these ruminants, as is observed endemically in Africa (*25*). Interestingly, alpaca, which are native to South America where OROV is endemic, are permissive to OROV infection (Fig. 1B, 2A, and 3E). Both RVFV and OROV are thought to use low-density lipoprotein receptor-related protein 1 (LRP1) as an entry receptor, and there is evidence that the cluster domains II and IV are particularly important for viral binding, whereas clusters I and III have not been shown to be involved(*26–28*). Sheep fibroblasts were strongly refractory to OROV, and thus we explored the role of LRP1 in OROV infection. Because OROV is re-emerging in South America, and new reassortant strains are more pathogenic in humans(*29, 30*), we also tested a more contemporary OROV strain. We found that both strains were unable to infect cow and sheep fibroblasts (Fig. 3F). Next, we transduced these sheep fibroblasts with cluster I or cluster IV from human LRP1, since cluster IV has previously been shown to bind OROV glycoproteins, whereas cluster I has not(*27*). We found that expression of cluster IV promoted infection in the presence or absence of IFN signaling while cluster I had a more limited effect, only revealing increased infection upon JAK inhibition (Fig. 3G, supplemental Fig. 3B-C). These data demonstrate that some ruminants are refractory to infection due to an entry defect that may be exacerbated by innate immune control.

### Conserved and differential responses to innate immune sensing and IFN signaling across species

We next evaluated the gene expression differences in antiviral sensing and signaling that may be driving virus resistance and susceptibility. We generated dynamic transcriptomes at baseline and using different stimuli. We treated with poly(I:C) to mimic RNA virus infection which induces pattern recognition receptor (PRR) signaling. This leads to activation of IRF3/IRF7 which induces IFNs and other innate factors. IFNs produced then induce JAK/STAT signaling and ISGs. We also directly treated with uIFN to evaluate ISGs in the absence of the upstream pathway. Lastly, we treated with poly(I:C) in the presence of JAKi to identify genes induced by RNA sensing alone in the absence of JAK/STAT signaling. Despite the rapid expansion in the availability of high-quality whole genomes(*31–33*), there are still large gaps in Mammalia. We therefore performed two parallel analyses; first, mapping to reference genomes where available; and second, performing *de novo* assembly and annotation to capture transcriptomes for species where no annotated reference genome is available. We benchmarked these approaches directly in species that have well-annotated reference genomes and found that both pipelines organize samples by species and treatment condition similarly (Supplementary Fig. 4A). Furthermore, analyzing all RNA-seq libraries by hierarchical clustering, we found strong correlations between replicates of the same species and treatment (median *r* = 0.994) (Supplementary Fig. 4B-C). Notably, correlations were higher between samples of the same species under different treatment conditions (median *r* = 0.914) than between different species under the same treatment (median *r* = 0.711), suggesting that species identity, rather than innate-immune stimulation, is the primary determinant of transcriptome state. We evaluated assembly quality by Benchmarking Universal Single-Copy Orthologs (BUSCO) ortholog completeness, and as expected found complete BUSCOs ranged from 71.8% to 78.2% (Supplementary Fig. 4D). For species with an annotated reference genome, we compared per-gene differential expression from both pipelines and found high concordance (cross-pipeline log2FC Pearson r, median 0.904; sign-agreement, median 85.8%; significant-DEG Jaccard index, median 0.703) and sufficient gene-symbol recovery across both pipelines (median 11,779 shared symbols per species) (Supplementary Fig. 4E-F). Together this generated 252 transcriptomes (63 species and 4 experimental conditions).

As expected, related species had similar gene expression profiles which deviated as evolutionary distance increased (Fig. 4A). We confirmed broad responsiveness to uIFN and poly(I:C) across species, with poly(I:C) inducing more genes than uIFN (Fig. 4B). Consistent with virus infection data (Fig. 2B) we found that cells from hedgehog, eastern cottontail rabbit, common marmoset and ring-tailed lemur were insensitive to uIFN treatment (Supplementary Fig. 4G). To determine if this was due to reduced responsiveness to IFN-I in general or incompatibility with uIFN, we compared gene expression response between species-specific IFN-I and uIFN in human, horse, pig, Jamaican fruit bat, eastern cottontail rabbit, domestic rabbit, and grey short-tailed opossum. We found that species-specific IFNβ and uIFN induce similar levels of MX1 in human, horse, pig, and Jamaican fruit bat, consistent with the RNAseq data and further validating the broad reactivity of uIFN (Supplementary Fig. 4H). We observed induction of MX1 and CXCL10 in rabbits and marsupials, respectively, upon treatment with species-specific IFN-I but not uIFN (Supplementary Fig. 4H). These data suggest that while uIFN can broadly activate IFN responses it does not signal equivalently across all mammals likely due to divergence of IFN receptors.

**Figure 4.**
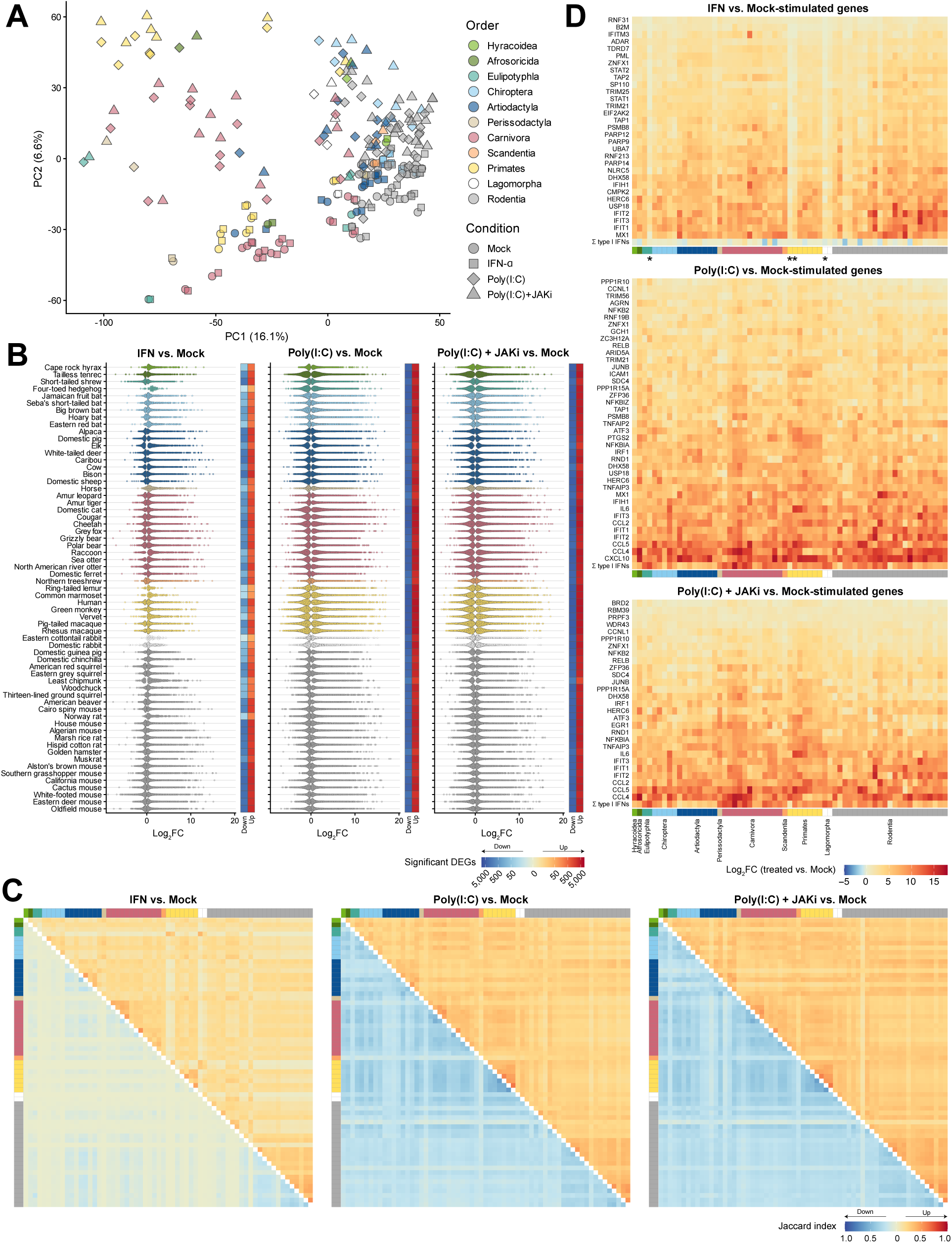
Comparative transcriptomics during virus sensing and IFN signaling in 63 unique species. (A) Principal components analysis of centered log2-scaled normalized counts across 63 species and four treatment conditions using *de novo* assembled transcriptomes. Each point is the centroid of the replicate PC scores for each species-condition combination (all replicates are shown in Supplementary Fig. 4A). (B) Distributions of significant differentially expressed genes (DEGs) across species. Each point shows a gene with an FDR < 0.05 in the indicated contrast and species. A point’s horizontal position shows that gene’s shrunken log2 fold change. Point color denotes mammalian order. Adjacent tiles show the number of significant downregulated and upregulated genes in blue and red, respectively. (C) Pairwise similarity of species-level DEG sets. Each species’ genes were classified as upregulated or downregulated using an FDR <0.05 and the sign of the shrunken log2 fold change. Tile color shows the pairwise Jaccard index (intersection over union) between each species’ DEG sets. The upper-right triangle shows overlap in upregulated DEG sets, and the lower-left triangle shows overlap among downregulated DEG sets. White diagonal tiles indicate self-comparisons. The margin color bars identify each species’ mammalian order. (D) Shrunken log2 fold changes for upregulated core DEGs in response to IFN (31 genes), Poly(I:C) (40 genes), and Poly(I:C) pretreated with JAKi (28 genes). A gene was defined as core when it was significant (FDR <0.05) and direction-consistent in all species. For the IFN-stimulated core-set calculation, Eastern cottontail rabbit, four-toed hedgehog, ring-tailed lemur, and common marmoset were excluded. These species are indicated with asterisks below the IFN vs. mock heatmap (see Supplementary Fig. 4G).

We observed gene expression differences with uIFN, poly(I:C), and poly(I:C) in the presence of JAKi with related species sharing gene expression signatures which diverged across species (Fig. 4C). We first focused on IFN treatment and by analyzing 59 species with robust responses to uIFN. We identified a core set of 31 ISGs showing induction across these species. Analysis across the full set show that they are shared across all species tested (Fig. 4D). 29 of these were in the set of 62 ISGs with conserved expression previously identified in a comparison of 10 vertebrate species(*9*). Our core ISGs are likely an underestimate as additional ISGs may be upregulated if stimulated with species-specific IFN-I. We also identified 40 core poly(I:C)-induced genes across all 63 species tested. Many of these genes overlap with those previously found in primates and rodents(*34, 35*). Pretreatment with JAKi prior to poly(I:C) stimulation revealed the PRR response independent from IFN signaling. This yielded 28 PRR-dependent genes shared across all species which include classic IRF3 targets IFIT1 and CXCL10(*36, 37*). At least one type I or III IFN subtype was induced in response to poly(I:C) or poly(I:C) in the presence of JAKi in all species, but no IFN subtype was shared across all species (Fig. 4D). Together these data demonstrate that there is a conserved transcriptional program downstream of virus sensing and signaling across mammals.

While many genes are co-regulated, we hypothesized that evolutionary trajectories would lead to some differential responses across species. First, we performed clade enrichment analyses, in which each gene’s change in expression within a given mammalian order was compared against the pooled response in all other orders. The difference in median fold-changes was then used to identify genes with clade-specific induction. Similar to previous reports, we found species- and order-specific ISGs, poly(I:C)- and PRR-stimulated genes (Fig. 5A)(*9*). We focused on genes involved in pathogen sensing and signaling and found that IRF3, traditionally not viewed as an ISG as it is not transcriptionally induced in humans or model organisms(*38*), is robustly induced in a subset of related artiodactyl species (Fig. 5B). We evaluated the promoter region of IRF3 for presence of IFN-stimulated response element (ISRE) and found several bovids but not cervids with canonical ISREs, suggesting a potential mechanism for IFN induction but also highlighting additional unknown factors at play (Fig. 5B). IRF3 induction by uIFN was validated by western blot demonstrating induction in cow and white-tailed deer but not human and alpaca, consistent with RNAseq data (Fig. 5C). Interestingly, a negative regulator of RIG-I signaling upstream of IRF3, RNF125 (*39*) was significantly induced by poly(I:C) in these species but not humans or commonly used model organisms, suggesting a potential regulatory mechanism compensating for the IFN induction of IRF3 (Fig. 4B). We also evaluated expression divergence at the family level and found that rodents in Cricetidae had significantly higher induction of cytokine signaling 1 (SOCS1) compared to other families in Rodentia as well as other orders (Supplementary Fig. 5A-B). Together these data demonstrate divergence in the response and regulation of seemingly canonical antiviral pathways across mammals.

**Figure 5.**
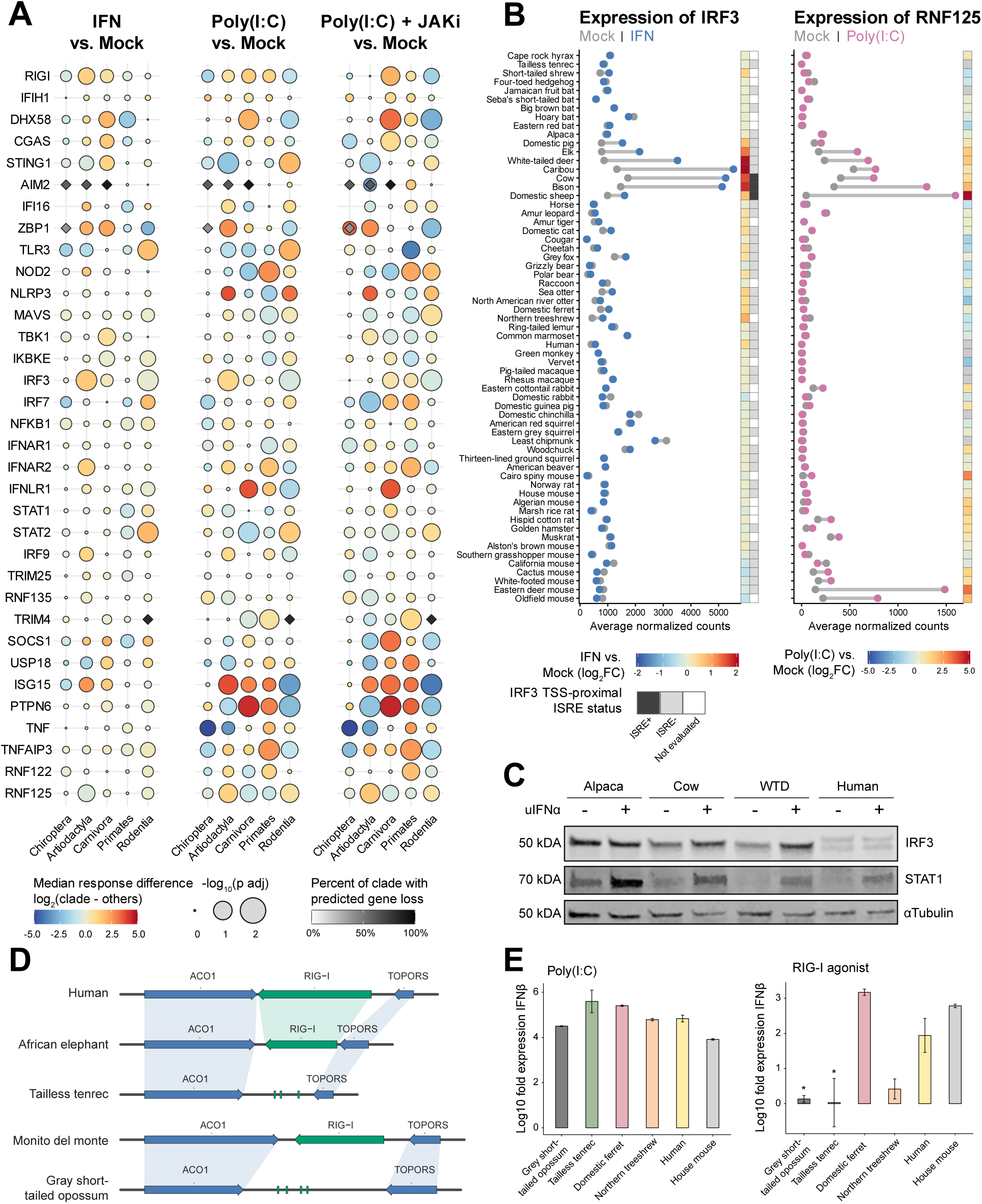
Differential regulation of antiviral sensing and signaling across mammals. (A) Order-level differences in innate immune gene responses under three treatment conditions. Log2 fold changes relative to mock were compared between species within each order versus all other species using two-sided Wilcoxon rank-sum tests. Bubble fill shows the focal order median minus the median across all other orders, and bubble size shows a -log10 p-value Benjamini-Hochberg adjusted across all genes detected in at least 32 of 63 cell lines. Diamonds mark predicted gene losses in at least two sampled species within an order and are shaded according to the percentage of species with a predicted loss. Genes are predicted as lost when at least two TOGA2 reference projections call a gene lost with no projections supporting an intact call. Only orders with at least five cell lines are shown. (B, left) IRF3 expression is IFN-stimulated in bovids and cervids. Each row shows a species’ DESeq2-normalized IRF3 counts under mock (grey) and universal IFN-α (blue) treatment. The left column of tiles shows the IFN-α versus mock log2FC values for each species. The right column of tiles shows the presence of a single shared regulatory locus near the IRF3 transcription start site. (B, right) RNF125 expression after poly(I:C) stimulation. Each row shows a species’ DESeq2-normalized RNF125 counts under mock (grey) and poly(I:C) (purple). The column of tiles shows the poly(I:C) versus mock log2FC values for each species. (C) The indicated species were treated with 1000 U/mL of uIFN and IRF3 and STAT1 were probed 24 hours after treatment. WTD = White-tailed deer. (E) Linear homology plots of the RIG-I (DDX58) locus in human (GRCh38.p14), African elephant (mLoxAfr1.hap2), tailless tenrec (mTenEca1.hap1), monito del monte (mDroGli1.pri), and the grey short-tailed opossum (mMonDom1.pri) reference assemblies. The tenrec locus contains no intact DDX58 gene, with three protein-level fragments recovered by tBLASTn. The grey short-tailed opossum locus also contains no intact DDX58 gene but retains sequences homologous to monito del monte coding exons 10, 14, 16 and 18. (E) The indicated species were treated with either poly(I:C) or RIG-I specific agonist and IFNβ levels were evaluated by qRT-PCR 24h post stimulation. Data are shown as log10 fold change over tubulin and untreated and are representative of 2 independent experiments.

In addition to changes in expression, we also evaluated loss of genes involved in innate immunity. When we analyzed core pathway components, we found loss of RIG-I in tailless tenrecs and in grey short-tailed opossum but not in related elephants and monito del monte opossums, respectively (Fig. 5D). Birds have also lost RIG-I and have concomitantly lost the RIG-I cofactor RIPLET(*40*). Chinese tree shrew have also lost RIG-I and it is possible our related northern tree shrew is also RIG-I deficient (*41*). Interestingly, using TOGA2 we also observed loss of the required RIG-I cofactor RIPLET, in northern tree shrews, grey short-tailed opossum and tailless tenrec. We further analyzed tenrecs and tree shrews where we stimulated the cells with mixed molecular weight poly(I:C) to trigger both RIG-I and MDA5 and found cells from tailless tenrecs and northern tree shrew had robust gene induction (Fig. 5E). Thus, Chinese tree shrews may have compensated for the loss of RIG-I significantly altering MDA5 to recognize a broader array of RNA(*41*). We tested for selection among mammalian MDA5 orthologues using adaptive branch-site random effects likelihood (*42*) and found that while there was evidence of episodic diversifying selection in the tree shrew lineage (p=0.015) there was no evidence of selection found for tailless tenrec (p=1). We therefore hypothesized that tenrec cells would be unable to respond to RIG-I agonist. Consistent with this hypothesis, grey short-tailed opossum and tailless tenrec had significantly lower induction of IFNβ in response to a RIG-I agonist compared to humans (Fig. 5E). Northern tree shrew also had reduced response to RIG-I stimulation (Fig. 5E). Together these data suggest significant unappreciated loss of antiviral sensors within mammals with the potential to impact susceptibility to virus infection.

### Host features impact susceptibility to infection

To determine if differential gene signatures can be linked to infection outcome, we combined our RNAseq and infectivity data. For each virus, we first determined how many species controlled replication via IFN signaling (i.e. no replication in untreated cells but replication in JAKi-treated cells, see methods), and how many did not control virus via IFN signaling (i.e. replication in untreated cells, see methods). Dengue virus (MOI = 1, 72h post-infection) exhibited a balance of species which do (N = 21) and do not control infection via IFN (N = 17), and we chose to further examine the relationship of gene expression and infection control for this virus.

We correlated gene expression levels under uIFN stimulation with percent infectivity for each species, and ranked genes based on their correlation coefficient. Negatively correlated genes could be putatively antiviral. However, we did not find canonical ISGs known to control DENV2 infection as top hits (Fig. 6A). We reasoned that this could be due in part to gene divergence and/or that combinations of genes may be necessary for control of infection. Gene divergence in STAT2 can provide a barrier to DENV infection (*19*). We therefore used rule-based modeling using R.ROSETTA(*43*) to identify combinations of genes and their expression values that can classify animals as either able or unable to control DENV2 infection via IFN signaling. We compared gene expression differences between untreated and uIFN-treated cells for the set of 38 animals with known viral control status. We constructed a rule-based model by first performing feature selection with Monte Carlo Feature Selection (*44, 45*) and generated the IF-THEN rules with R.ROSETTA(*43*) using a list of ISGs which we defined as genes upregulated between uIFN treatment and mock in at least 10% of species (padj < 0.05 and log2FC > 1) (Supplementary Table 6). The model achieved strong performance at classifying species by DENV2 control status (accuracy = 0.9188, AUC = 0.9815). Combinatorial IF-THEN rules generated by the model revealed co-predictive networks(*46*) that were centered on IRF8, REC8 and ATAD1 which control antiviral responses (Fig. 6B and Supplementary Fig. 6A). These genes positively regulate antiviral sensing and signaling and may promote a more effective antiviral state(*47, 48*). Importantly OASL, which is antiviral against DENV2 was identified in the network as a combinatorial regulator with REC8 (Fig. 6B, Supplementary Table 6)(*49*). Additionally, FBXO39 and IRF8 have inverted roles in the networks with FBX039 absent in controlled group and up in non-controlled. Interestingly this gene plays a proviral role during RVFV infection (*50*). These genes would not have been identified in traditional correlation analyses (Supplementary Fig. 6B) as they are not highly anticorrelated with DENV2 infection. These data demonstrate that gene expression across species can identify potential antiviral programs that control infection outcomes.

**Figure 6.**
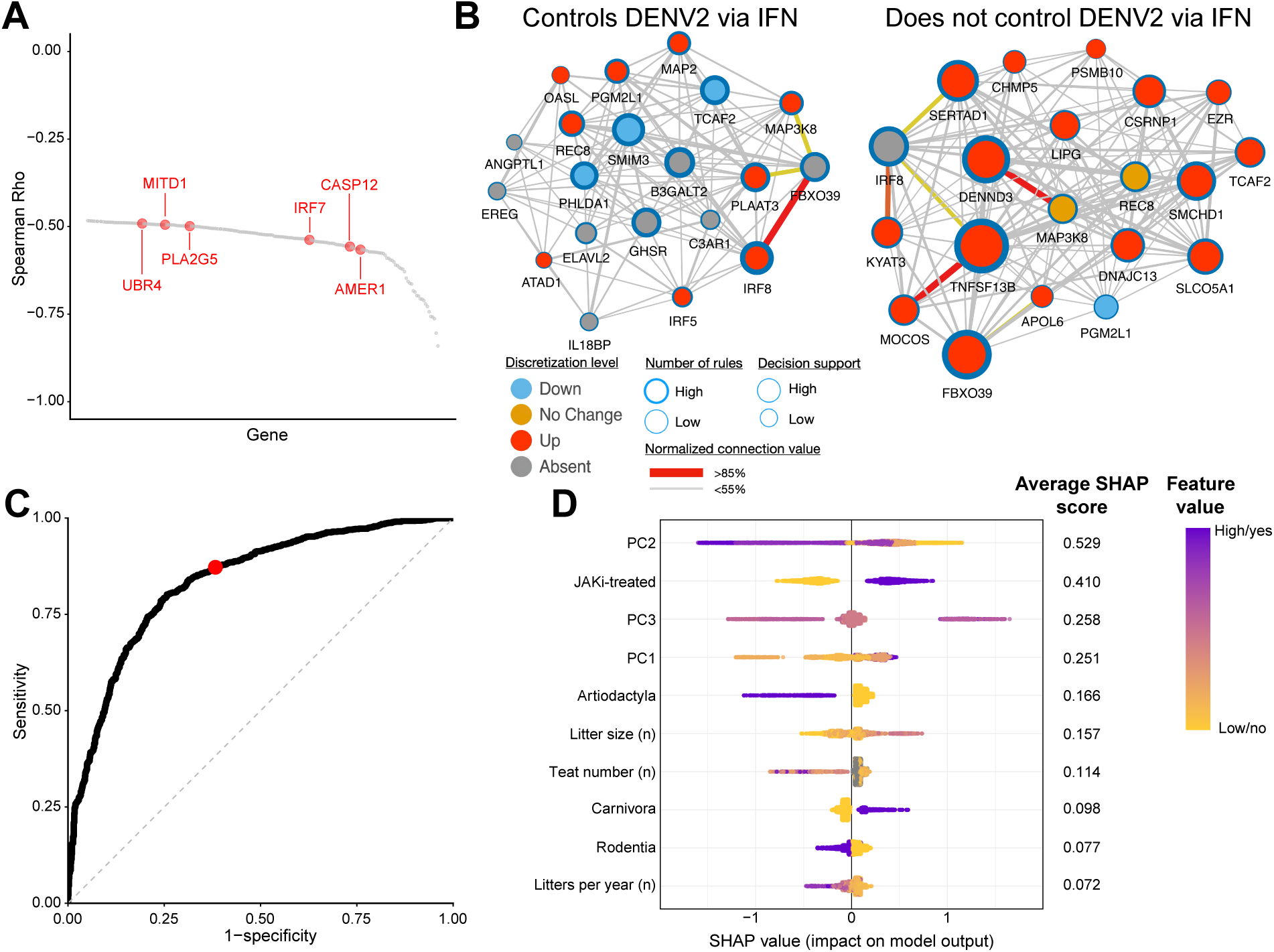
Interpreting viral infection outcomes using machine learning. (A) Rank plot of correlation coefficients of infection outcomes in untreated fibroblasts versus gene expression levels in uIFN-treated cells. Depicted are the top 200 genes whose expression levels (normalized TPM) are significantly anti-correlated with % infection (FDR < 0.05). Red dots highlight uIFN-stimulated genes present in at least 10% of mammalian species profiled with RNAseq. (B) Rule-based model decision network of genes associated with control of DENV2 via IFN (left) or lack of IFN-mediated control of DENV2 (right). (C) ROC AUC of XGBoost model of virus-host compatibility comparing true positive versus false positive rates. Red dot indicates the sensitivity and specificity for a model with a probability cutoff of 0.5 chosen for distinguishing positive versus negative infection outcomes. (D) SHAP beeswarm plot of top 10 features impacting model output. SHAP values describe how much each predictor influenced the predicted infection outcome for individual virus-host pairs. Each dot per predictor row is a virus-host pair. The color of the dot represents the value of the predictor for the individual virus-host pair. The position on the X axis indicates how the value of that predictor relates to the outcome. Positive SHAP values indicate that the predictor’s value is related to a higher probability of infection in that virus-host pair.

In addition to gene expression and regulation, virus and host traits may be associated with infection outcome in unexpected ways. We used machine learning to determine whether host and/or viral features could predict replication compatibility. We compiled 32 features describing viral genome structure, mammalian life history traits from the COMBINE database(*51*), and cell IFN signaling status. These were then used to fit an extreme gradient boosting (XGBoost) model which predicted presence or absence of viral replication in both untreated and JAKi-treated cells. Including the JAKi-treated data led to greater instances of viral replication compared to non-replication. The fitted model achieved an ROC AUC of 0.88, 89% sensitivity, 67% specificity, and precision of 83% and 81% accuracy (Fig. 6C). The model was fit using ten-fold cross-validation stratified by host species. A second model was fit using ten folds stratified by virus species which achieved comparable performance metrics (ROC AUC = 0.84, sensitivity = 87%, specificity = 62%, precision = 80%, and accuracy = 78%).

We calculated SHAP values of the predictors (Fig. 6D) and found that codon usage biases (transformed into PCs), JAKi-treatment, Artiodactyla order, and litter-related life history traits were the most importance predictors of virus-host compatibility. Viruses including RVFV and orthoflaviviruses which less frequently use GGA, TGG, and CCG codons (associated with PC2) and more frequently use GAA, GAG and ACA codons (associated with PC3) were more likely to infect a greater range of mammalian hosts (Fig. 6D, Supplementary Fig. 6C-D). Artiodactyla mammals were less likely to be infected by the viruses in our panel. Litters per year and litter size had highest importance among the life history traits examined. We calculated the marginal effect of feature value on infection probability and found that features consistently impacted outcome in the same direction across both untreated and JAKi-treated cells (Supplemental Fig. 6E). For example, higher litter size and more litters per year were associated with greater probability of infection regardless of cell treatment. This suggests that IFN responses are not underlying these relationships. Taken together, this shows life history traits and viral genomic features can predict probability of viral replication in fibroblasts, however the high sensitivity and low specificity suggest the model was more effective at identifying virus-host pairs with high compatibility than low compatibility.

## Discussion

The study of virus-host interactions has been dominated by a few model organisms. Given the substantial evolutionary pressure placed on organisms by pathogens there is likely a wealth of antiviral mechanisms that control infections in the wild that have yet to be discovered. These may predict host compatibility and virus evolutionary trajectories and ultimately be repurposed for new interventions. To evaluate antiviral responses across mammals we built a library of primary dermal fibroblasts from animals that have never been experimentally studied. These may be key reservoirs for viruses and/or have evolved novel mechanisms to restrict infection. This approach relies on live cells which can be used to directly address molecular barriers (e.g. entry, innate immunity, egress, etc.) and can evaluate complete antiviral sensing and signaling pathways. This has allowed us to report a dynamic data set of virus-host interactions with 15,867 pairwise observations and 252 transcriptomes.

Our data reveal that innate immunity is a major barrier across viruses and hosts and is particularly evident across orthoflaviviruses. For example, DENV specificity for degrading human STAT2 poses a barrier in mice which are poorly permissive due to this incompatibility; this may be more general across orthoflaviviruses and hosts(*19*). Conversely, Rift Valley fever virus displays high infectivity even in the face of IFN responses consistent with NSs targeting a highly conserved component of the core RNA pol II machinery(*22, 23*). RVFV is not thought to broadly infect animals in Africa suggesting additional factors are underlying its limited spread *in vivo* compared to our *in vitro* system. VEEV also demonstrated low sensitivity to antiviral responses across species suggesting it may have a more generalist strategy for innate immune evasion and antagonism.

While each virus can infect some species, we find that 22% of virus-host pairs show low infectivity at high MOIs and upon inhibition of JAK-STAT signaling. This suggests an incompatibility with one or more host factors required for viral infection. Since we are monitoring infection by viral antigen production, the block can be at multiple stages of replication including entry and virus transcription and translation. All the viruses used in this study can infect humans and we observed infection of human fibroblasts in all cases. Importantly, these data are consistent with studies that found that many viral glycoproteins can enter human cells *ex vivo*, suggesting that entry may be a weak barrier(*52*). These generalist arboviruses can replicate in arthropod and mammalian hosts, and it is thus not surprising that there is widespread infection in general across the fibroblast zoo. However, there specific examples where there is an incompatibility even in the absence of antiviral; signaling which reveals additional host factors that are essential for infection. For example, several orthoflaviruses across bat species. We provide examples that highlight clear differences in entry across species where we find that CHIKV infection of bats is under complex control. Likewise, we reveal a strong entry defect in ruminants for OROV infection.

Viral antagonism of innate defenses has driven selection and diversification across antiviral sensing and signaling pathways. Nevertheless, there is a small core of ISGs induced in response to virus sensing and IFN signaling across Mammalia, consistent with previous observations(*9, 34, 35*). We also identified genes involved in distinct steps in innate defenses including PRRs, PRR and IFN signaling pathway genes, and ISG regulators that are fundamentally different across Mammalia. RIG-I has been independently lost in multiple lineages, and may also have been lost in Cetaceans, star nosed mole, cape golden mole, tree shrew, tenrec and short-tailed opossum(*41, 53, 54*). It is likely that differential sensing drives the infection phenotypes we observed across the zoo. Few viruses infect artiodactyls at baseline, but many are controlled by IFN signaling. Whether this is due to altered regulation of IRF3 or other innate immune pathway components is an open question.

The experimental virus-host data generated here is particularly useful for training machine learning models given that it is non-sparse and highly dynamic, unlike other training datasets which may rely on surveillance data and are less effective at identifying cases where hosts are resistant to viruses. Our preliminary modeling highlighted several potential features underpinning higher probability of virus replication, including several host features which may have co-evolved in response to high pathogen burden (e.g. higher litter sizes and fewer litters per year). We also identified codon usage as a key feature correlating with virus replication probability which is in line with previous studies examining human-infecting viruses(*55*). However, while we have strong diversity of mammals represented in the data, the comparatively low diversity of viruses may hinder ranking of virus features that are important for replication across mammals. As we include increased virus diversity and we suggest that there will be a more balanced data set with positive and negative replication outcomes in the future, which we anticipate will both improve our understanding of virus-host interactions and the performance of machine learning models aimed at predicting virus-host compatibility.

While these datasets provide valuable insights into virus-host interactions across mammals there are several limitations. These are primary cells which have the advantage of intact innate immune systems which are often perturbed in transformed cell lines. However, primary cells grow at different rates making it difficult to make MOIs identical across species. Major phenotypes reproduce across experiments and individuals (Supplementary Fig. 1E-F), but smaller differences will need to be independently verified. Currently our data from most species are only represented by a single individual. There could be significant genetic diversity within species which is not captured here. This study only reports data from a single cell type which captures approximately 65% of the transcriptome of a given species(*56*). Fibroblasts are highly relevant for arboviruses, and all the viruses used in this study infect human fibroblasts to some extent. However, fibroblasts are not the natural cellular tropism of all viruses. Future studies can validate and extend these findings to additional cell types induced from fibroblasts which have been differentiated into iPCS(*57–59*). Comparative transcriptomic studies are hampered by lack of genomes and/or quality of annotation. We attempted to overcome this using *de novo* assembly and BLAST-based annotation to identify transcripts. However, a limitation of this approach is that it has difficulty distinguishing between paralogs, which is particularly important for innate effectors as many known antiviral genes are products of gene duplications(*60–62*). The rapid progress in genome sequencing, assembly, and identification of orthologs will alleviate this concern over time and will permit direct mapping to genomes of diverse species(*31–33, 63*). “Universal” IFN is a chimera of two IFNα’s with documented broad reactivity across Mammalia but cannot activate ISGs in avians(*64*). However, our results demonstrate that uIFN does not signal equivalently across all species and therefore conserved responses and absence of induced ISGs should be interpreted with caution. Using species-specific IFN across all these species may not be feasible. However, poly(I:C) is not species-specific and demonstrates core and unique responses some of which overlap with uIFN. Despite these limitations we believe these data sets have covered important biology of virus-host interactions which will spark new research directions.

## Materials and Methods

### Fibroblast generation and characterization

Primary dermal fibroblast cultures were generated using skin tissue sourced from surgical biopsies or necropsies. To extract cells, tissue was digested for 1 hour in Liberase TM (Roche) and, the following day, separated from debris using a 2.5mM EDTA solution and a 100um cell strainer (Corning). Each culture was expanded and maintained at 37°C and 5% CO_2_ in complete DMEM, supplemental antibiotics, non-essential amino acids (Thermo Fisher), and 55uM 2-mercapto-ethanol. For experiments, fibroblast lines were utilized between passages 3 and 10. Species identity was verified through CO1 barcode sequencing(*65*). A small group of the primary cell lines required different conditions for stable growth; that and other information including tissue source and sex is included in Supplementary Table 1.

### Virus infections

Information on viruses used in experiments, including strains, stock growth method, stock quantification, and MOIs, is included in Supplementary Table 2. Prior to each infection, primary cells were plated and pretreated with 1000U/mL or 1200U/mL universal interferon alpha (uIFN, PBL) and 10μM ruxolitinib (Selleck) for 1 or 2h and 1h, respectively. Virus was diluted in serum-free DMEM, was added at a high and low MOI as indicated. Complete media was added after an hour; vehicle, uIFN and ruxolitinib conditions were reestablished. Virus levels were quantified at virus specific time points outlined in Supplementary Table 2.

### Virus quantification by High-Throughput Imaging

Viral infectivity assays were done in 96-well black-walled clear-bottom plates (Corning). Plates were washed in DPBS with Ca^++^/Mg^++^ and fixed in 4% PFA or directly fixed with 8% PFA (final concentration 4%). Cells were permeabilized using 0.1% Triton-X/PBS solution and blocked with 5% normal goat serum, 3.5% BSA, and 0.05% Tween20 in PBS. Some cells were blocked with PBS with 2% Bovine Serum Albumin (BSA) and 0.1% Triton-X100. Viral proteins were immunolabelled with primary antibody overnight and a goat anti-IgG H+L Alexa Flour (Invitrogen) secondary for 2 hours (antibody list included in Supplementary Table 2). Nuclei were dyed with a Hoechst 33342 Solution.

Using the BioTek/Agilent Cytation 5 Cell Imaging Multi-mode Reader or automated imaging (ImageXpress, 10x objective, two fluorescent channels, 9 images per well) with automated image analysis using MetaXpress, each infection condition and corresponding replicate were captured with a 2x2-tiled image using the 10X objective. Light settings for DAPI (377, 447), TRIT-C (556, 600), and CY5 (628, 685) channels were optimized to each virus, but kept the same across cell lines, time points, and experiments. Images were preprocessed in the BioTek Gen5 Image Prime 3.15 software package (v.3.15.15.0) using the default rolling ball algorithm settings. Infectivity was measured by identifying total cell count using Hoechst-positive signal and quantifying number of cells associated with viral protein signal using a distance-based secondary mask. Raw cell counts were exported to Microsoft Excel 2016 (v.16.0.4266.1001). Imaging methods differed slightly for some viral infections due to equipment availability between collaborating labs.

All data was combined across experiments and assessed for quality; failed image replicates were removed. Ruxolitinib and uIFN fold-change analyses were calculated as follows:

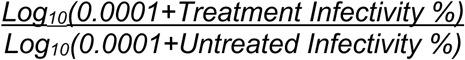

### Lentivirus generation and fibroblast transduction

To generate lentiviral particles, cells were seeded in 10cm dishes at 5x10^6^ cells per dish. The next day, cells were transfected with the plasmid of interest, psPAX2, and pMD2.G at a 2:2:1 mass ratio with LipoD293 (SignaGen SL100668). MXRA8 plasmids and LRP1 cluster plasmids were a gift courtesy of Dr. Michael Diamond, Washington University in St. Louis. Cells were incubated for 18 hours, then the medium was replaced with fresh complete DMEM. Viral supernatants were then collected 24 hours later, spun at 1,200 rpm for 5 minutes, and filtered through a 0.45-µm filter Viral particles were concentrated using the PEG Virus Precipitation Kit (Abcam, ab102538), following the manufacturer’s instructions. Fibroblasts were seeded in 6-well plates, and 50uL lentivirus encoding mouse MXRA8, empty vector, human LRP1 cluster I, or LRP1 cluster IV were added to cells. MXRA8-expressing fibroblasts were selected with 5ug/mL blasticidin. LRP1 cluster-expressing fibroblasts were selected with 1ug puromycin.

### Live-cell imaging

Immediately post infection with CHIKV-mKate, cells in a 96-well plate were added to the Incucyte SX1 live-cell analysis instrument. Cells were imaged every 2 hours for 88 hours. Red image mean (mKate signal) was measured to detect infection over time.

### Cell treatment and RNA library preparation for sequencing

Cells were transfected with 1µg/mL of a 1:1 mixture of high- and low-molecular weight poly(I:C) (Invitrogen) using Lipofectamine 2000 (Invitrogen), both with and without a one-hour 10μM ruxolitinib (Selleck) pretreatment. In parallel, cell lines were treated with 1,000U/mL of universal IFN-alpha (PBL). Six hours after treatments, cells were lysed and RNA was extracted using the MagMAX mirVana Total RNA Isolation Kit (Applied Biosystems). RNA was quantified using Quant-it RiboGreen (Invitrogen) and quality tested using an Agilent TapeStation. Libraries were prepared using the mRNA library prep kit from Watchmaker Genomics and sequenced on the AVITI24 system (Element Biosciences) to generate paired-end 150-bp reads at the University of Minnesota Genomics Center.

### Read mapping to annotated reference genomes

RNA-seq reads from species with an annotated genome assembly (Supplementary Table S3) were processed using the nf-core/rnaseq pipeline v3.24.0(*66*). Read quality was assessed using FastQC v0.12.1 before and after trimming with Trim Galore v0.6.10 and Cutadapt v5.2 (*67*) which removed adapter sequences, trimmed 3’ bases with Phred scores below 20, and discarded read pairs when either read was shorter than 20 bp. Trimmed reads were aligned to the corresponding reference genome using STAR v2.7.11b(*68*). Transcript abundances were quantified with Salmon v1.10.3. Unmapped reads were screened for contaminants with Kraken2 v2.1.6 (*69*) and Bracken v3.0.1 (*70*) against the PlusPF database. Differential expression was modeled separately for each cell line with DESeq2 v1.50.2(*71*), retaining genes with at least 10 counts in at least 3 samples. Where several genes shared a symbol, the most highly expressed one represented that symbol. Wald p values were adjusted with the Benjamini-Hochberg procedure, and log2 fold changes were shrunk using ashr v2.2-63(*72*).

### De novo transcriptome assembly and annotation

RNA-seq reads from all species were processed using the transXpress Snakemake pipeline(*73*). Read quality was assessed using FastQC v0.12.1 before and after trimming with Trimmomatic v0.39(*74*), which removed adapter sequences, trimmed leading and trailing bases with Phred scores below 5, truncated reads where the mean quality in a 4-base sliding window fell below 5, and discarded read pairs when either read was shorter than 25 bp. Trimmed reads from all samples of a cell line were pooled and de novo assembled with Trinity v2.13.2 (*75*) using default parameters. Assembly completeness was assessed using BUSCO v6.0.0 against the mammalia_odb10 dataset (*76*) (Supplementary Fig. 3B). Assembled contigs were annotated by blastx searches using NCBI BLAST+ v2.15.0 (*77*) against the UniProtKB/Swiss-Prot database (release 2025_04)(*78*). Only contigs with a UniProt gene name and a blastx E-value below 1E-6 were retained. Hits to human proteins were assigned HGNC symbols (January 6, 2026 release). Hits to non-human proteins were assigned the human gene name listed by NCBI Gene as the one-to-one ortholog (as of September 2, 2026). All other hits retained their UniProt gene names. Transcript abundances were quantified with kallisto v0.48.0(*79*). Counts for transcripts assigned the same gene name were summed. Differential expression was modeled separately for each cell line with DESeq2 v1.50.2(*71*), retaining genes with at least 10 counts in at least 3 samples (Supplementary Table S4). Wald p values were adjusted with the Benjamini-Hochberg procedure, and log2 fold changes were shrunk using ashr v2.2-63(*72*).

### Western blots

Equal protein quantities from lysates were diluted 1:1 in Laemmli buffer (Bio-Rad) containing 2-Mercaptoethanol (0.28M) and boiled at 95°C for 5 minutes. Samples were run at RT in Mini-Protean® TGX™ Precast gel with either a 4-15% or 4-20% gradient (Bio-Rad) and a Precision Plus Protein Dual Color Standard (Bio-Rad). Blots were transferred to 0.45µM nitrocellulose at 4°C in 1x Transfer Buffer (Bio-Rad) containing 20% methanol. Blots were blocked for 30 minutes at RT with 5% Dry Milk Powder in TBS-T (10 mM Tris base pH 7.5, 150mM NaCl, 0.1% Tween-20, dH_2_O ad 1L) and incubated overnight at 4°C on a shaker with primary antibody. The following day, blots were washed with TBS-T, incubated in 1:1000 secondary antibody solution for 1 hour and imaged with the LI-COR Odyssey Fc Imager. Primary rabbit (rb) *a*’IRF3 (gift from Michael Gale Jr) and rabbit (rb) *a*’STAT1 (T5168, Sigma) were used at 1:1000 and mouse (ms) *a*’Tubulin (9175, Cell signaling) at 1:2000. Secondary antibodies IRDye® 680LT Donkey *a*’ms IgG IRDye® 800CW Donkey *a*’rb IgG (LI-COR biosciences) were used at 1:1000.

### qRT-PCR for species-specific gene expression

To assess RIG-I detection, cells were transfected for 24 hours as previously described, with 1ug/mL poly(I:C) in parallel with 400ng/mL of 5’ppp dsRNA RIG-I agonist (Invivogen). Also, to test universal IFN non-responders, cells were treated with 50ng/mL and 200ng/mL of species-specific IFN-β and 13ng/mL uIFN for 6 hours. RNA was harvested using MagMAX mirVana Total RNA Isolation Kit (Applied Biosystems). After RNA was quantified with Quant-it RiboGreen (Invitrogen), cDNA was generated from 250ng of sample using the High Capacity Reverse Transcription Kit (Applied Biosystems). Gene expression was assessed using SYBR Select Master Mix (Applied Biosystems) and the QuantStudio 5 PCR System (Applied Biosystems). Cycle threshold was set using the default settings in the QuantStudio Design and Analysis Software (v1.5.3). Gene expression, averaged between three biological replicates, was normalized to an untreated control and pan-mammalian housekeeping gene. Primer sets are included in Supplementary Table 5.

### Rule based modeling

Monte Carlo Feature Selection (MCFS) generated a list of top-ranked genes (version 1.3.6). We performed additional refinement with the stepwise selection method also called sequential forward selection method to find an optimal set of genes that classify a species as being able or unable to control DENV2 via IFN. Rule-based models were built using 5, 10, 15, … 150 ranked genes and assessed by applying 4-fold cross-validation. The model that achieved the highest accuracy during this evaluation was designated as the final one. In case of ties, we evaluated the models based on the least number of genes used. Rule-based modeling was conducted using the ‘rosetta’ function in the R library -R.ROSETTA (version 2.3.4.1)(*43*), which implements rough set theory(*80–82*). The parameters used by the ‘rosetta’ function in each run were as follows: the reducerDiscernibility was set to ‘Object’ with ‘Controlled by IFN’ as the fallback class and ‘Genetic’ as the reducer type. Rule filtration was enabled with ruleFilterAccuracy set to 80% and ruleFilterSupport values based on percentages of observation of 5%.

The visualization of rules was generated using the ‘visunetcyto’ function from the csVisuNet R library (version 1.4.2.4)(*46*), which is specifically designed for visualizing rule-based networks in Cytoscape. This function accepts input from the ‘rosetta’ function in the R.ROSETTA library (version 2.2.9), which generates a set of IF-THEN rules to differentiate between decision classes (outcomes). The IF-part of a rule consists of a single descriptor (for example, a gene and its expression and discretization level) or, more generally, of a conjunction of descriptors. For instance, IF IRF8 = “Up” AND FBXO39 = “Absent” THEN Decision = “Controlled by IFN” and IF IRF8 = “Absent” AND FBXO39 = “Up” THEN Decision = “Not controlled by IFN”. The “IF” part denotes a combination of descriptors that predict the decision class. Significantly, rules for each decision class are visualized. Each rule can be evaluated using quality metrics such as accuracy, support, and decision coverage. Using a set of rules, csVisuNet constructs a rule network. The nodes of the network are formed by the descriptors and each pair of descriptors occurring in a rule creates an edge of the network. The thickness of an edge is determined by:

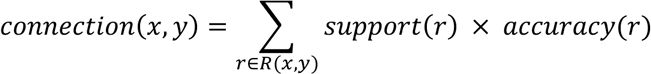

where *R(x,y)* is the set of all rules containing *x* and *y* in the model.

The color of the edge is defined to represent the strength of connection(*83*): red indicates a strong connection (normalized connection value ≥ 85%), yellow represents a medium connection (normalized connection value ≥ 55% and < 85%) and grey signifies a weak connection (normalized connection value < 55%). The size of a node indicates the mean support. The color of the node represents the expression or discretization level (Down, No Change, Up, as defined in our project), while the node border indicates the number of rules associated with that descriptor. Once a set of rules is transcribed into a rule network, it enables the identification of interdependencies and co-predictive factors related to a decision. It also highlights hubs, similarly to hubs in computer networks; in this context, a hub forms numerous connections to other descriptors and is crucial for distinguishing the decision class.

The csVisuNet (version 1.3.9) library employs a ‘visuArc’ function(*84*) that highlights hub genes and their connections within the network. It employs a specialized visualization known as Arc Representation, which illustrates the arcs connecting genes directly to a selected hub gene, irrespective of their expected behaviors within the gene network. Arc diagrams can be generated based on a specified decision class, allowing for a focused analysis of the network in relation to a chosen outcome. It is important to note that the rule networks show associations of the outcome and not any causal relation. However, these associations may possibly serve as hypotheses of causality to be used for experiments to confirm or reject them (*85*).

### Gradient boosting

We applied a gradient boosting machine learning model to assess how host and virus traits influenced the likelihood of successful infection. Features for machine learning were compiled for the viruses and hosts depicted in Figure 1. A set of 55 host features and 12 virus features was initially chosen. Host features (N=54) were compiled from the COMBINE database(*51*). Mammals listed in Figure 1 without data in COMBINE were dropped from further analysis, resulting in 77 hosts available for model fitting. A host feature describing mammalian order was also included for a total of 55 features. The virus features included a single feature each describing genome sense, whether the virus is segmented, number of proteins, number of genes, genome length, combined CDS (coding sequence) lengths, GC frequency, and virus taxonomic family. Four additional features were included to describe codon usage bias of open reading frames. These four features were determined by calculating virus codon usage frequencies of the in-frame nucleotide sequences of all open reading frames in the viral genome using the coRdon package in R(*86*). Then, PCA was used to distill the 64 resulting frequencies into principal components (Supplementary Fig. 6C). The scree plot of the explained variance indicated that the first four PCs explained the majority of variability and thus these four PCs were included as viral genome features. The model was fit on both untreated and JAKi-treated data, and therefore a final variable was included to describe the JAKi treatment status of the virus-host infection data.

Categorical features were one-hot encoded using the mltools package in R (https://doi.org/10.32614/CRAN.package.mltools), yielding an initial set of 95 features. Highly co-correlated feature pairs were flagged by high Spearman correlation coefficients (absolute ρ greater than 0.7). Among these pairs, we chose to keep the more biologically relevant feature as a predictor or the feature with higher complete data across the COMBINE database. This yielded a final predictor set of 56 features.

Machine learning was used for binary classification of presence or absence of viral infection in a virus-host pair. A cutoff of 7% infection was used to distinguish between uninfected and infected cells. This cutoff was chosen by comparing the range of background infectivity in mock-infected cell lines, where available (mean = 1.46, standard deviation = 3.65). We assume completely resistant cells should have measured percent infectivity below 1.46% but chose a higher cutoff to ensure we are only considering positive infection in cases where there is a confident signal.

Ten-fold cross-validation with hyperparameter tuning was used to fit an XGBoost model using the xgboost package in R, version 3.4.2(*87*). Hyperparameters of learning rate (0.01-0.3), maximum tree depth (2–10), subsampling rate (0.5-1), column subsample ratio (0.5-1), L1 and L2 regularization terms (0–5), and minimum loss reduction (0–5) were fit using a random grid search targeting the highest ROC AUC. To prevent data leakage between training and test sets, folds were created by stratifying the data by either virus or host to separately fit two models. The host-and virus-stratified models were later compared for consistency based on performance metrics. The stratification was performed such that no host species (or virus species, for the virus-stratified model) was included in multiple folds. Folds were generated to maintain as much balance between positive and negative infection outcomes as possible. Tuned hyperparameters were used in a final cross-validation using the custom folds and 50 early stopping rounds. The out-of-fold predictions were generated during final cross-validation using a probability threshold of 0.5 to distinguish positive versus negative cases. These predictions were used to generate the ROC, confusion matrix and performance metrics using the caret packet in R(*88*). SHAP (SHapley Additive exPlanations) values were calculated using a final model trained on the full dataset with tuned hyperparameters using the SHAPforxgboost package in R(*89*). Marginal effects of feature values were calculated for each feature using the final trained model and data subset by JAKi treatment status with the stats package and viewed as partial dependence plots using ggplot2(*90*). Contributions of codon usage frequencies to the principal components was visualized using the factoextra package in R(*91*).

## Acknowledgements

We sincerely thank all the tissue donors listed in Supplementary Table 1. We also wish to thank the University of Minnesota Genomics Center for help with library preparation and sequencing. We wish to thank members of the Cherry lab and the UPENN High-throughput Screening Core (RRID SCD_022379).

## Data, code, and materials availability

RNA sequencing data are available as FASTQ files in the NCBI sequence read archive under BioProject no TBD. RNAseq count data are available under GEO accession number TBD. Code for the manuscript figures and analyses is available at https://github.com/langloislab/TBD. The primary cells are a finite resource, are not immortalized and are restricted by passage number. Amount of tissue donated and expansion rate in culture dictates the number of cells available. Some cell lines may not be shared as they are owned by the donating zoos and agency. We will make every effort to share cell lines where possible on request.

