## Supplemental figures for "A mammalian fibroblast zoo reveals barriers to cross-species virus replication and differential regulation of antiviral immunity"

### Supplementary figures

**A**

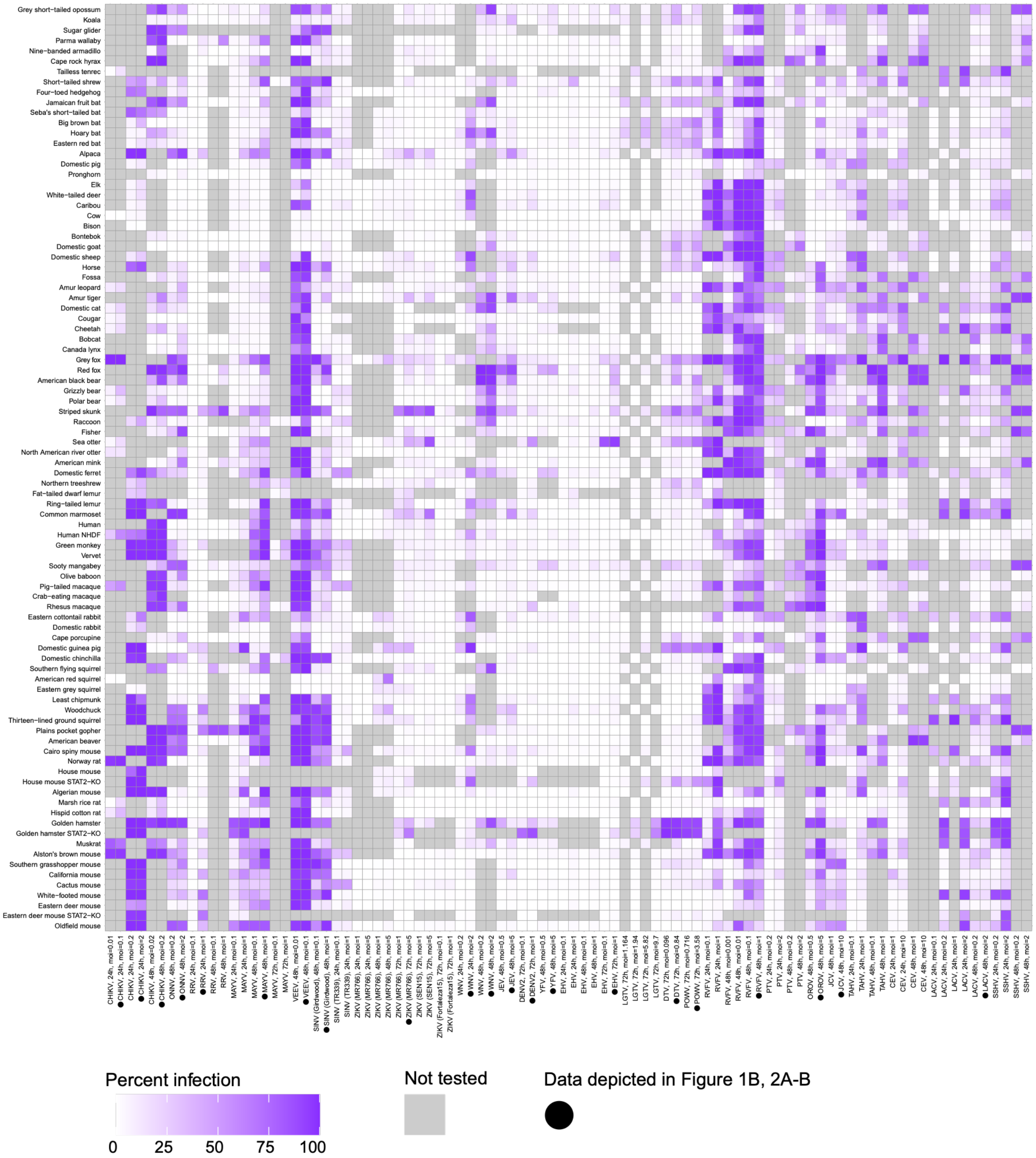

B

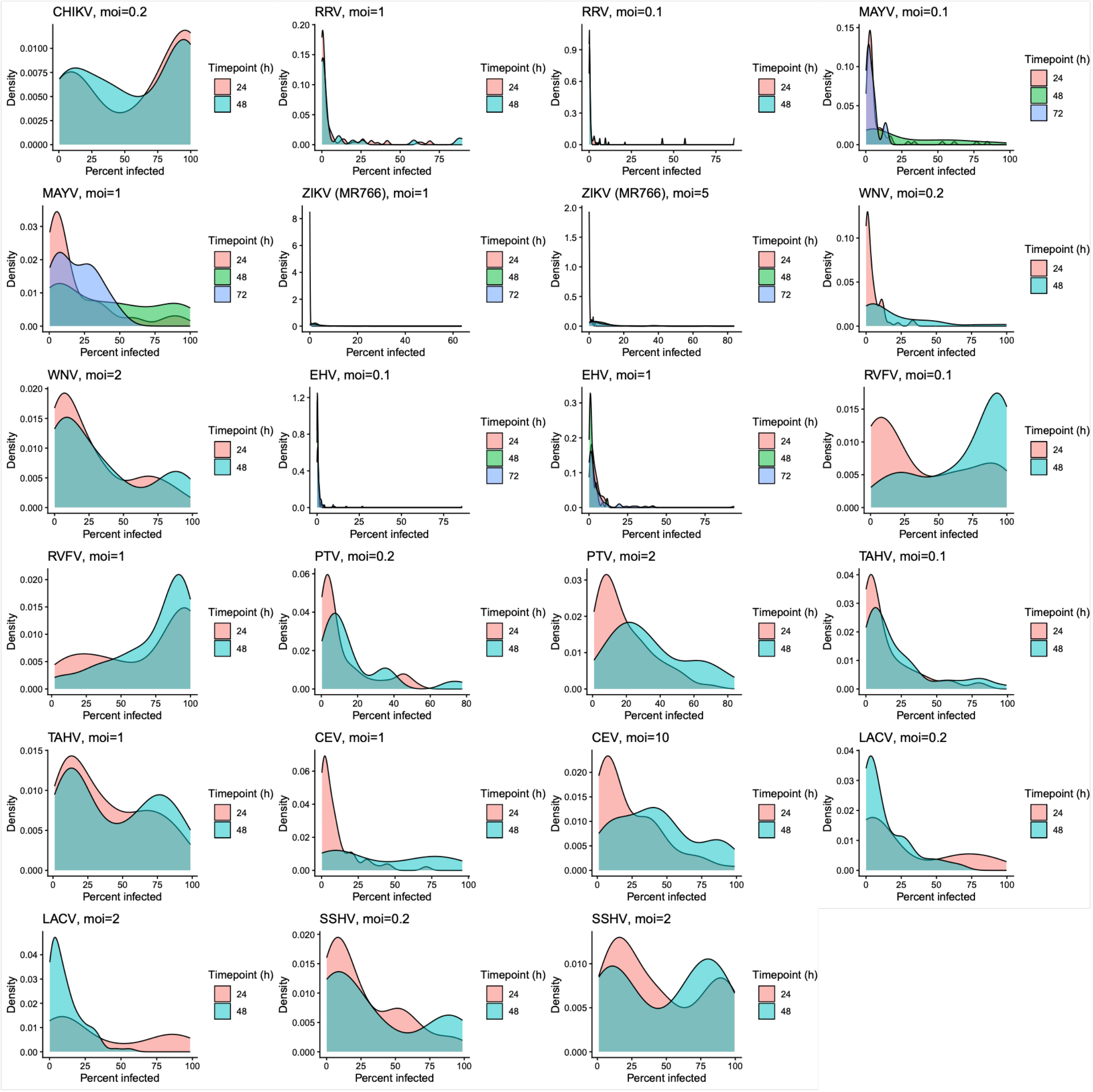
CHIKV, moi=0.2
RRV, moi=1
RRV, moi=0.1
MAYV, moi=0.1
MAYV, moi=1
ZIKV (MR766), moi=1
ZIKV (MR766), moi=5
WNV, moi=0.2
WNV, moi=2
EHV, moi=0.1
EHV, moi=1
RVFV, moi=0.1
RVFV, moi=1
PTV, moi=0.2
PTV, moi=2
TAHV, moi=0.1
TAHV, moi=1
CEV, moi=1
CEV, moi=10
LACV, moi=0.2
LACV, moi=2
SSHV, moi=0.2
SSHV, moi=2
Density
Percent infected
Timepoint (h)
24
48
72

C

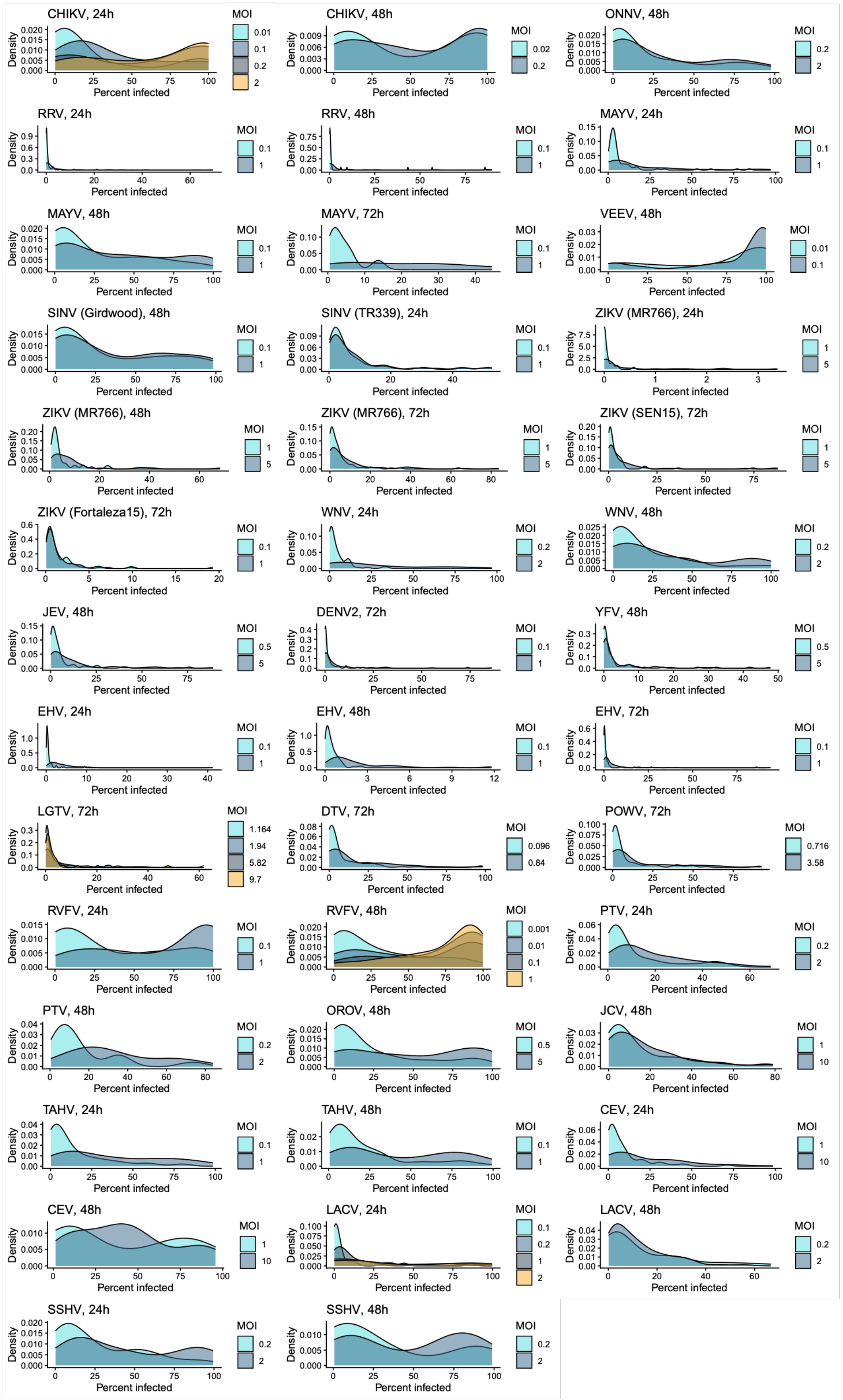
CHIKV, 24h
MOI 0.01 0.1 0.2 2
CHIKV, 48h
MOI 0.02 0.2
ONNV, 48h
MOI 0.2 2
RRV, 24h
MOI 0.1 1
RRV, 48h
MOI 0.1 1
MAYV, 24h
MOI 0.1 1
MAYV, 48h
MOI 0.1 1
MAYV, 72h
MOI 0.1 1
VEEV, 48h
MOI 0.01 0.1
SINV (Girdwood), 48h
MOI 0.1 1
SINV (TR339), 24h
MOI 0.1 1
ZIKV (MR766), 24h
MOI 1 5
ZIKV (MR766), 48h
MOI 1 5
ZIKV (MR766), 72h
MOI 1 5
ZIKV (SEN15), 72h
MOI 1 5
ZIKV (Fortaleza15), 72h
MOI 0.1 1
WNV, 24h
MOI 0.2 2
WNV, 48h
MOI 0.2 2
JEV, 48h
MOI 0.5 5
DENV2, 72h
MOI 0.1 1
YFV, 48h
MOI 0.5 5
EHV, 24h
MOI 0.1 1
EHV, 48h
MOI 0.1 1
EHV, 72h
MOI 0.1 1
LGTV, 72h
MOI 1.164 1.94 5.82 9.7
DTV, 72h
MOI 0.096 0.84
POWV, 72h
MOI 0.716 3.58
RVFV, 24h
MOI 0.1 1
RVFV, 48h
MOI 0.001 0.01 0.1 1
PTV, 24h
MOI 0.2 2
PTV, 48h
MOI 0.2 2
OROV, 48h
MOI 0.5 5
JCV, 48h
MOI 1 10
TAHV, 24h
MOI 0.1 1
TAHV, 48h
MOI 0.1 1
CEV, 24h
MOI 1 10
CEV, 48h
MOI 1 10
LACV, 24h
MOI 0.1 0.2 1 2
LACV, 48h
MOI 0.2 2
SSHV, 24h
MOI 0.2 2
SSHV, 48h
MOI 0.2 2
Density
Percent infected

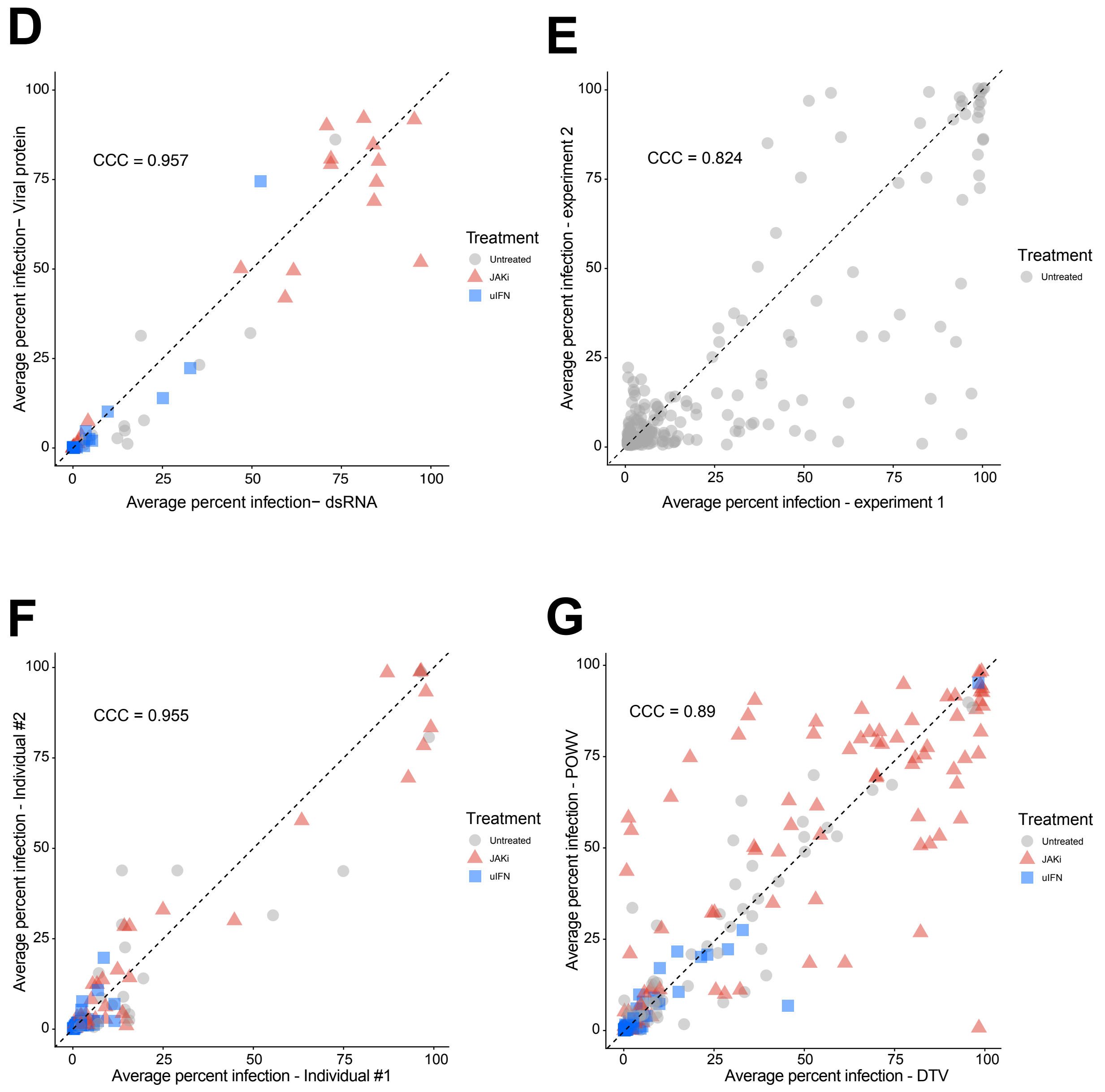


**Supplementary Figure 1. Pairwise comparisons of virus-host replication dynamics across the fibroblast zoo, related to Fig. 1.**

(A) Heatmap of all virus-host pair tested across the zoo with MOI and timepoints denoted on the X axis. MOI and timepoint combinations that are shown in in Fig. 1 are indicated with black dots. (B) Density plots of percent infections across time courses for select viruses and species indicated in panel A. (C) Density plots of percent infection across MOI ranges for select viruses and species indicated in panel A. (D) Concordance in percent infection as measured by antibody staining against virus protein vs. anti-dsRNA antibody staining. (E) Concordance of percent infection for species which were tested in multiple independent infection experiments. (F) Concordance of percent infection for genetically diverse individuals within species. Each dot represents a mammalian species and is colored by infection condition. X and Y-axis values represent percent infection for individuals 1 and 2. Hosts and viruses for D-F are indicated in Table S1 and S2, respectively. (G) Concordance of percent infection across genetically related flaviviruses, deer tick virus (DTV) and Powassan virus (POWV). Strain information listed in Table S1.

A

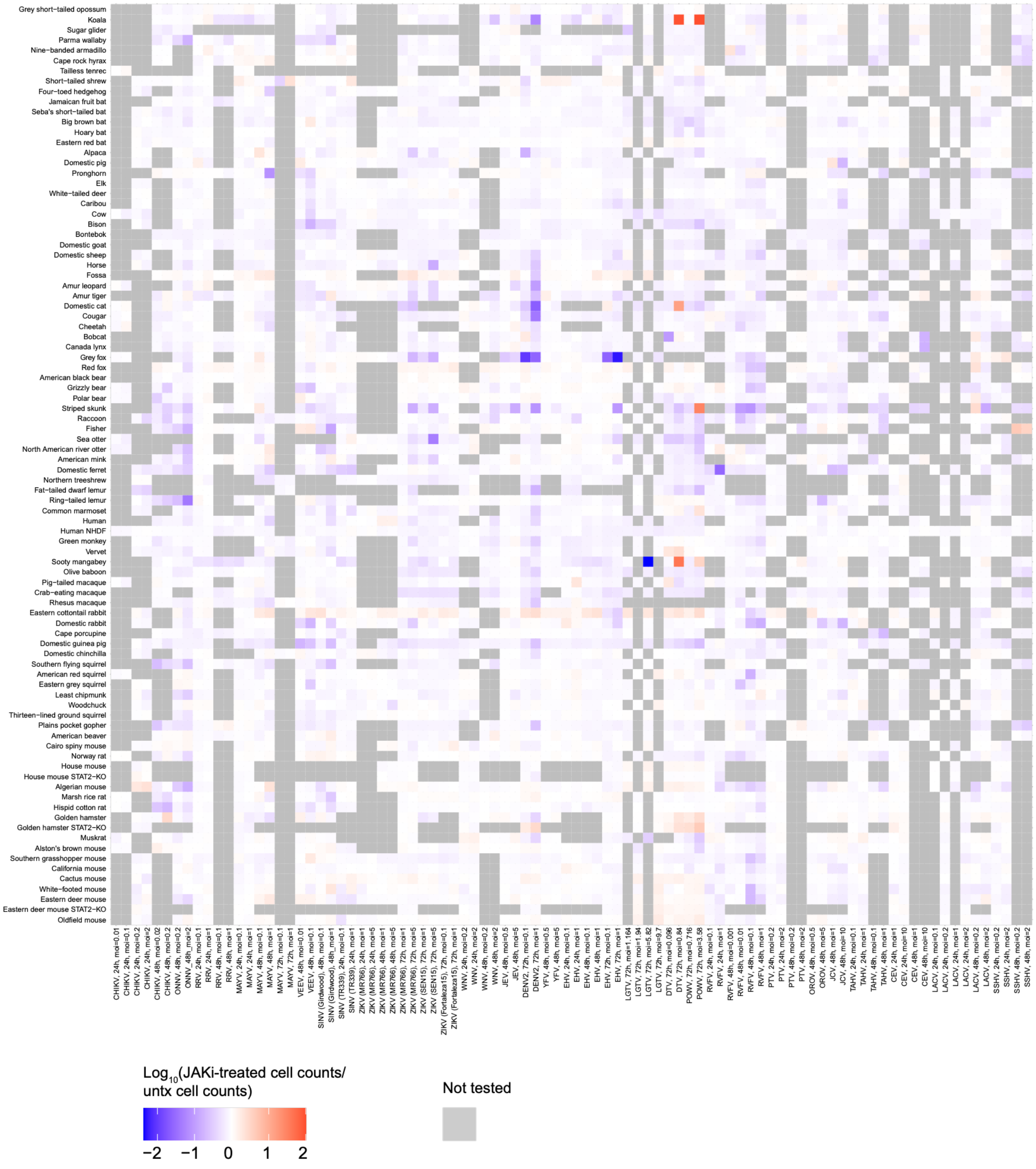
Grey short-tailed opossum
Koala
Sugar glider
Parma wallaby
Nine-banded armadillo
Cape rock hyrax
Tailless tenrec
Short-tailed shrew
Four-toed hedgehog
Jamaican fruit bat
Seba's short-tailed bat
Big brown bat
Hoary bat
Eastern red bat
Alpaca
Domestic pig
Pronghorn
Elk
White-tailed deer
Caribou
Cow
Bison
Bontebok
Domestic goat
Domestic sheep
Horse
Fossa
Amur leopard
Amur tiger
Domestic cat
Cougar
Cheetah
Bobcat
Canada lynx
Grey fox
Red fox
American black bear
Grizzly bear
Polar bear
Striped skunk
Raccoon
Fisher
Sea otter
North American river otter
American mink
Domestic ferret
Northern treeshrew
Fat-tailed dwarf lemur
Ring-tailed lemur
Common marmoset
Human
Human NHDF
Green monkey
Vervet
Sooty mangabey
Olive baboon
Pig-tailed macaque
Crab-eating macaque
Rhesus macaque
Eastern cottontail rabbit
Domestic rabbit
Cape porcupine
Domestic guinea pig
Domestic chinchilla
Southern flying squirrel
American red squirrel
Eastern grey squirrel
Least chipmunk
Woodchuck
Thirteen-lined ground squirrel
Plains pocket gopher
American beaver
Cairo spiny mouse
Norway rat
House mouse
House mouse STAT2-KO
Algerian mouse
Marsh rice rat
Hispid cotton rat
Golden hamster
Golden hamster STAT2-KO
Muskrat
Alston's brown mouse
Southern grasshopper mouse
California mouse
Cactus mouse
White-footed mouse
Eastern deer mouse
Eastern deer mouse STAT2-KO
Oldfield mouse
Log10(JAKi-treated cell counts/ untx cell counts)
−2
−1
0
1
2
Not tested

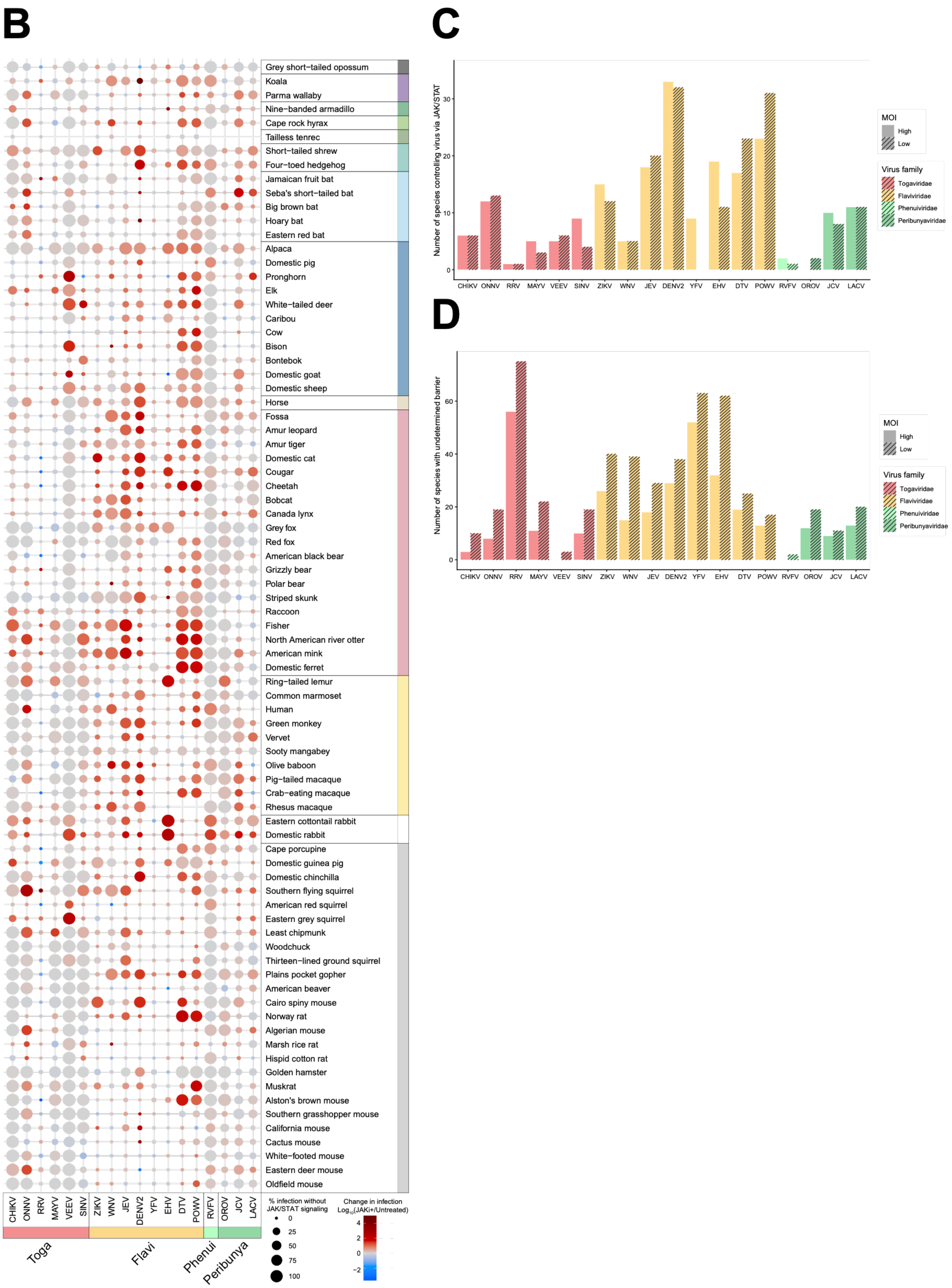


**Supplementary Figure 2. Evaluation of IFN as a barrier to infection across species, related to Fig. 2.**

(A) Heatmap of log10 fold change in cell counts of JAKi-treated compared to untreated condition for each virus-host pair. Grey boxes indicated that those viruses-host pairs were not tested or that cell count data is not available for comparison. (B) Cells from 81 species were pretreated for 1hr with JAKi or with media alone (untreated) and then infected at a 10-fold lower MOI than Fig. 2A. Cells were analyzed 24-72h later by immunostaining with virus-specific antigen. Percent infection with JAKi (size of dot) was determined by imaging. Log10 fold change in percent infection of JAKi-treated cells over untreated cells is shown as color of the dot (rescue of infection in red). See Table S2 for MOI time point and strain for each virus. (C) Comparison of number of species controlled by JAK/STAT signaling for each virus at high and low MOI (10-fold range). A mammalian species was considered to control a virus via JAK/STAT signaling if the percent infection in untreated cells was less than or equal to 5% and the infection in JAKi-treated cells was greater than 5% and there was at least a 0.7 log10 fold change between the JAKi-treated and untreated percent infection values. (D) Comparison of the number of species with an undetermined molecular barrier to infection. A barrier to infection could not be determined if viral infection was not detected under both untreated and JAKi-treated conditions.

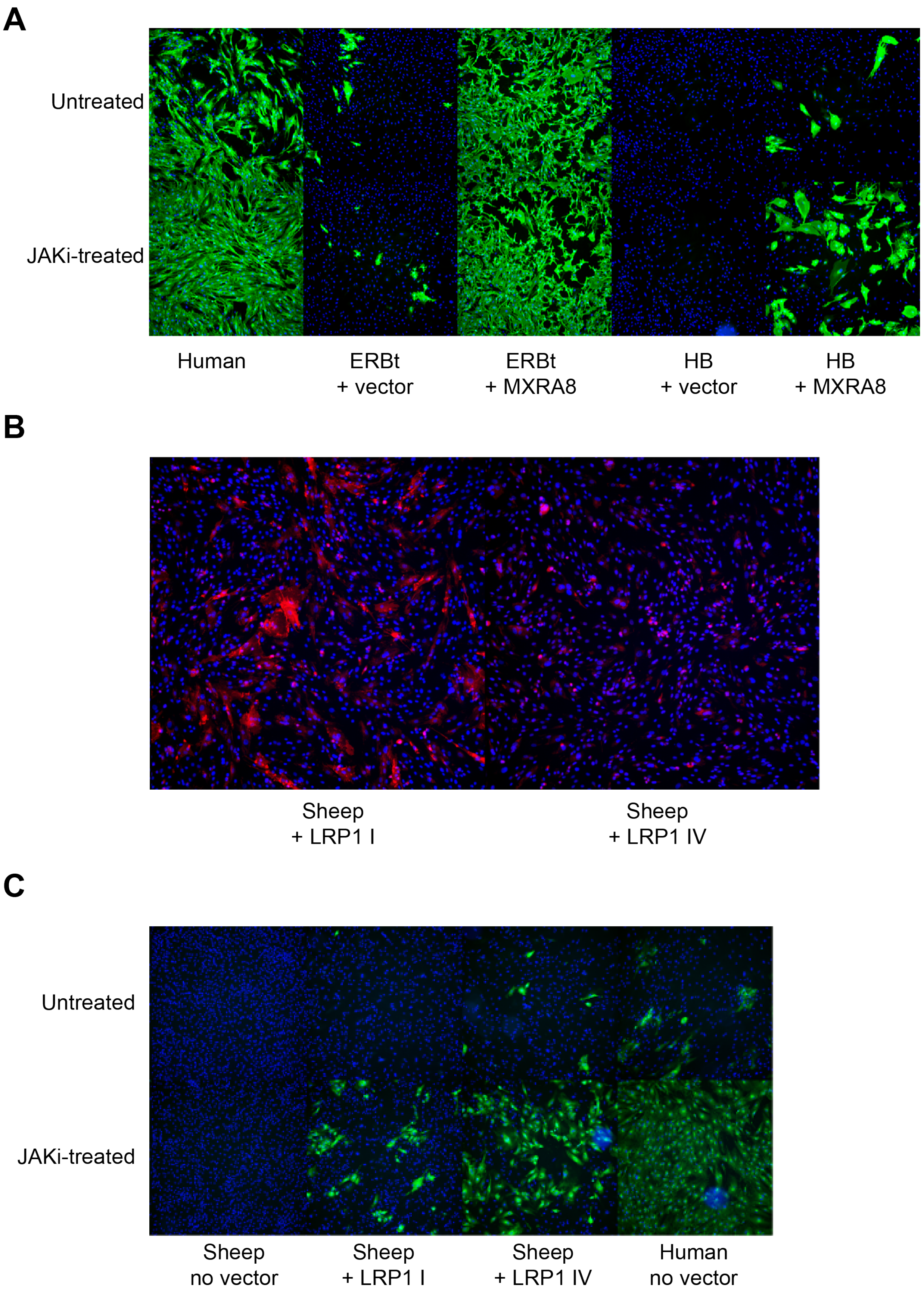
A
Untreated
JAKi-treated
Human
ERBt
+ vector
ERBt
+ MXRA8
HB
+ vector
HB
+ MXRA8
B
Sheep
+ LRP1 I
Sheep
+ LRP1 IV
C
Untreated
JAKi-treated
Sheep
no vector
Sheep
+ LRP1 I
Sheep
+ LRP1 IV
Human
no vector

**Supplementary Figure 3. Virus entry, related to Fig. 3.** (A) Eastern red bat and Hoary bat cells were transduced with mMXRA8 and selected with blasticidin and then were pretreated with DMSO, or ruxolitinib and infected with CHIKV (MOI 0.01) and fixed 24hpi. Infection was measured by immunofluorescence (IF) staining for viral antigen. (B) Sheep cells transduced with human LRP1 cluster I or cluster IV and selected with puromycin were stained for FLAG and imaged. (C) Human cells and sheep cells from (B) were pretreated with DMSO or ruxolitinib, infected with OROV (MOI 5) and fixed 48hpi. Infection was measured by immunofluorescence (IF) staining for viral antigen.

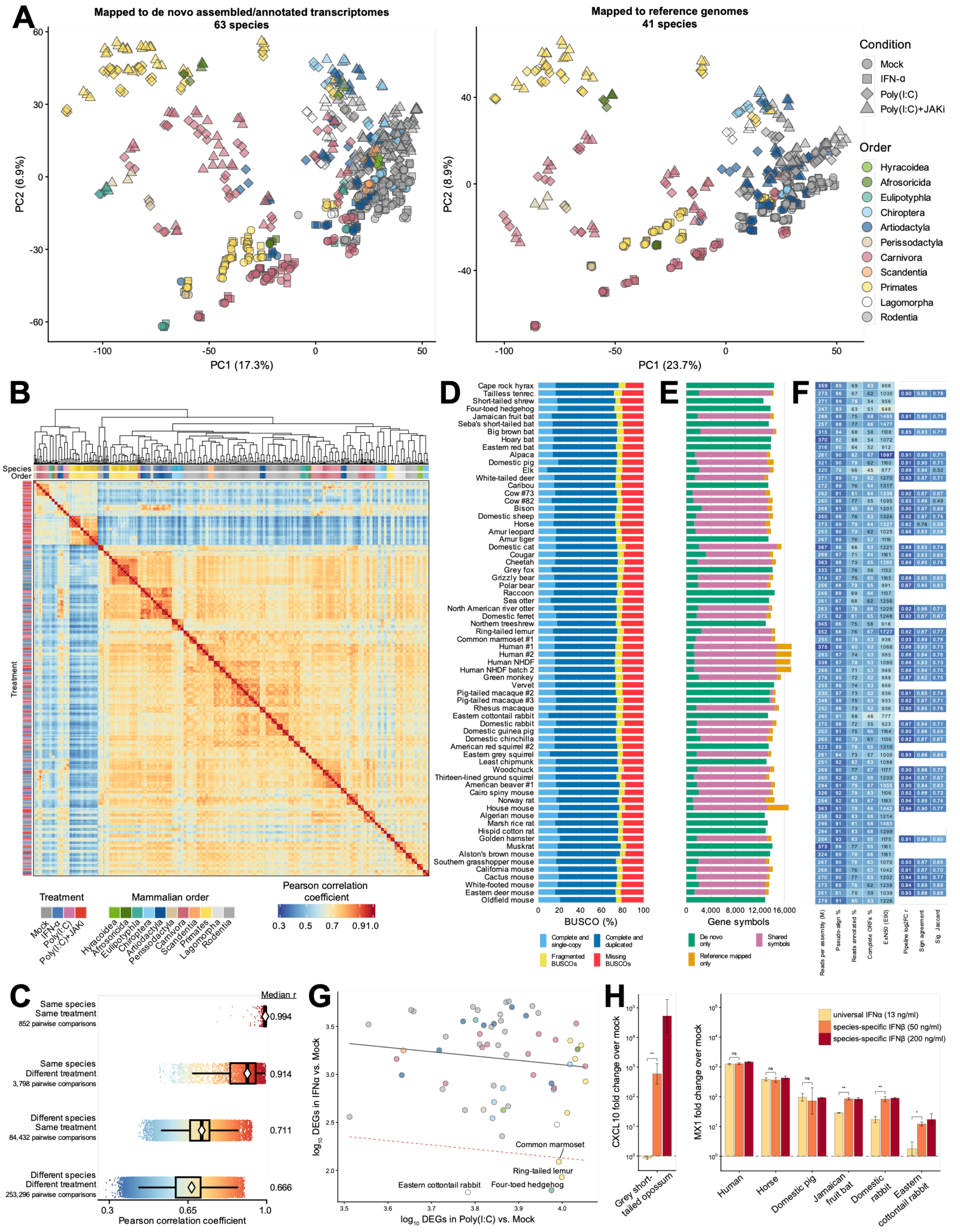


**Supplementary Figure 4. Transcriptome quality control and species-specific IFN responsiveness, related to Fig. 4.** (A) Principal components analysis of the two quantification pipelines, with all replicates shown. Left, *de novo* assembled and annotated transcriptomes:

principal components were computed using log2-scaled normalized counts from 9,521 gene symbols measured in every sample with non-zero variance. Right, reference genome-mapped data: the same procedure applied to 6,873 gene symbols shared across all reference-annotated samples. (B) Hierarchical clustering of 828 RNA-seq libraries, which clustered primarily by species. Pairwise Pearson correlations were computed across 9,521 shared, expression-filtered gene symbols using variance-stabilized expression from the *de novo* dataset. Samples were clustered by complete linkage using the distance 1-r. (C) Distribution of transcriptome-wide similarity among all 342,378 unique sample pairs from (B), partitioned into four classes. Boxes show the interquartile range, and diamonds mark the median. (D) BUSCO ortholog completeness of each *de novo* assembly against the mammalia_odb10 lineage. Complete BUSCOs ranged from 71.8% to 78.2% (median 74.4%). (E) Gene symbol recovery by the two pipelines. Stacked bars split each species' detected genes into symbols recovered by only one pipeline, or both. LOC-prefixed annotations are excluded from the reference-mapped set. (F) Per-species assembly and cross-pipeline quality metrics. The all-species block shows the total processed fragments contributing to each *de novo* assembly, the median percentage pseudo-aligned to the assembly, the median percentage of input fragments assigned to annotated transcripts, the percentage of TransDecoder-predicted ORFs classified as complete, and ExN50 at E90. For species with reference-mapped results, the second block shows the median across six treatment contrasts of the Pearson correlation between log2FC values, the fraction of shared symbols with concordant log2FC signs, and the Jaccard index of the significant-gene sets at FDR < 0.05. (G) Identification of species with a deficient response to universal IFN-α. Each point represents one species and plots its IFN-α versus mock DEG count (y-axis) against its Poly(I:C) versus mock DEG count (x-axis). DEGs were counted from the *de novo* dataset at FDR < 0.05 in either direction. The grey line shows the regression fit across all species; the red dashed line is the fit minus two median absolute deviations. (H) Species-specific IFN-β induces ISGs more strongly than universal IFN-α in some species. CXCL10 or MX1 induction measured by qRT-PCR, normalized to α-tubulin and expressed as fold change ($2^{-\Delta\Delta Ct}$) over each species' own mock. Cells were treated with universal IFN-α (13 ng/ml), or with species-specific IFN-β at 50 or 200 ng/ml in biological triplicate. Welch t-tests compare universal IFN-α with species-specific IFN-β at 50 ng/ml within each species (*P < 0.05; **P < 0.005).

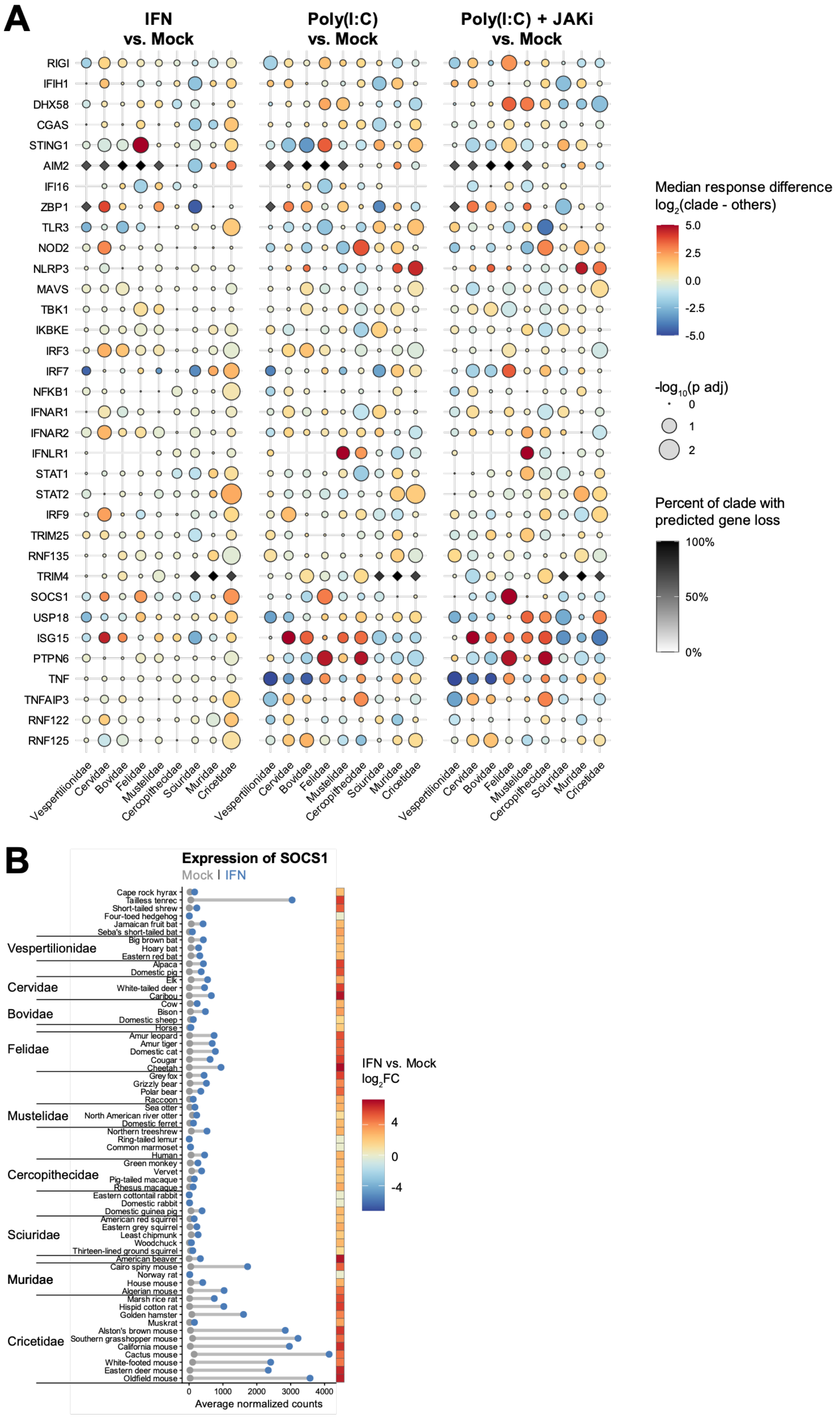


**Supplementary Figure 5. Divergent antiviral responses across mammals, related to Fig. 5.**

(A) Family-level differences in innate immune gene responses under three treatment conditions. Log2 fold changes relative to mock were compared between species within each family and all other species using two-sided Wilcoxon rank-sum tests. Bubble fill shows the focal family median minus the median across all other families, and bubble size shows a -log10 p-value Benjamini-Hochberg adjusted across all genes detected in at least 32 of 63 cell lines. Diamonds mark predicted gene losses in at least two sampled species within a species and are shaded according to the percentage of species with a predicted loss. Genes are predicted as lost when at least two TOGA2 reference projections call a gene lost with no projections supporting an intact call. Only families with at least five cell lines are shown. (B, left) SOCS2 expression is IFN-stimulated in cricetidae and several other species. Each row shows a species' DESeq2-normalized SOCS2 counts under mock (blue) and universal IFN-α (red) treatment. The left column of tiles shows the IFN-α versus mock log2FC values for each species.

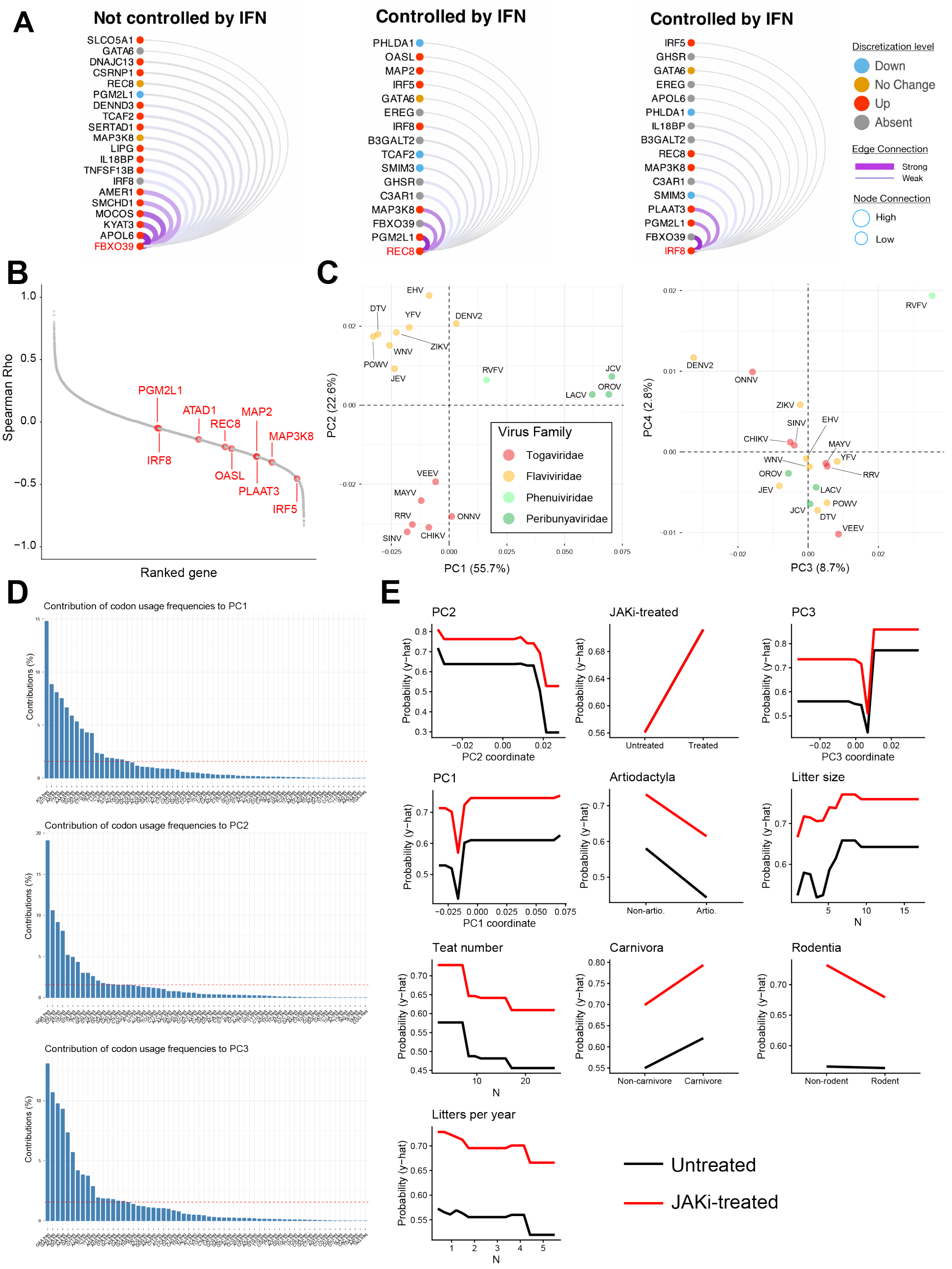
A
Not controlled by IFN
Controlled by IFN
Controlled by IFN
Discretization level
Down
No Change
Up
Absent
Edge Connection
Strong
Weak
Node Connection
High
Low
B
Spearman Rho
Ranked gene
PGM2L1
ATAD1
REC8
MAP2
MAP3K8
IRF8
OASL
PLAAT3
IRF5
C
PC1 (55.7%)
PC2 (22.6%)
PC3 (8.7%)
PC4 (2.8%)
Virus Family
Togaviridae
Flaviviridae
Phenuiviridae
Peribunyaviridae
D
Contribution of codon usage frequencies to PC1
Contribution of codon usage frequencies to PC2
Contribution of codon usage frequencies to PC3
Contributions (%)
E
PC2
JAKi-treated
PC3
PC1
Artiodactyla
Litter size
Teat number
Carnivora
Rodentia
Litters per year
Probability (y-hat)
Untreated
JAKi-treated

**Supplemental figure 6. Alternative extreme gradient boosting model of virus-host compatibility and codon usage PCA, related to Fig. 6.** (A) VisuArc's of the Central Nodes in DENV2 R.ROSETTA model (genes REC8, IRF8 and FBXO39). Subtitle under the gene name indicates whether relationships are derived from the decision networks associated with DENV2 control by IFN (i.e. REC8 and IRF8) or lack of DENV2 control by IFN (i.e. FBXO39). (B) Univariate correlation of infection outcomes in untreated fibroblasts versus gene expression levels in uIFN-treated cells. Each dot represents a gene, ranked by the Spearman Rho correlation of expression levels under uIFN stimulation (normalized TPM) versus percent infection (FDR < 0.05). Red genes are genes whose upregulation was identified by R.ROSETTA decision network for DENV2 control by uIFN (i.e. red nodes in Fig. 6B, left). (C) PCA plots of codon usage frequencies used by viruses in the XGBoost training data. Top plot shows PC1 versus PC2, bottom plot shows PC3 versus PC4. (D) Contribution of codon usage frequencies to variance explained by PC1, PC2 and PC3. (E) Partial dependence plots of the top 10 features based on average SHAP score. Dependence plots show the impact of feature value (x-axis) on probability of virus replication (y-axis), for both the untreated (black) and JAKi-treated (red) datasets.

**Supplementary Table 1.** Species and meta data for the fibroblast zoo

**Supplementary Table 2.** Viruses and metadata for infection studies

**Supplementary Table 3.** Reference genome information for fibroblast zoo species

**Supplementary Table 4.** De novo assembly methods and results summary

**Supplementary Table 5.** qPCR primers

**Supplementary Table 6.** R.ROSETTA IF-THEN rules
